# Human CCR4-NOT suppresses pervasive transcription and retrotransposable elements

**DOI:** 10.64898/2026.01.16.699749

**Authors:** Shardul Kulkarni, Alexis Morrissey, Aswathy Sebastian, Oluwasegun T. Akinniyi, Cheryl A. Keller, Istvan Albert, Shaun Mahony, Joseph C. Reese

## Abstract

CCR4-NOT regulates multiple steps in gene regulation and has been well studied in budding yeast. Although primarily cytoplasmic, where it plays an essential role in mRNA degradation, the human complex has poorly characterized nuclear functions. Here, we used auxin-induced degradation to rapidly deplete the scaffold subunit CNOT1 and the E3 ligase CNOT4, and characterized the transcriptional functions of the human CCR4-NOT complex. Using transient transcriptome profiling (TT-Seq) to measure ongoing transcription, we found widespread activation of RNA synthesis in depleted cells across genic and intergenic regions. Interestingly, fewer genes were repressed, including KRAB-Zinc-Finger-protein (KZNF) genes, especially those on chromosome 19. KZNFs repress genes and retrotransposable elements (rTEs), and consistent with decreased KZNF expression, rTEs, mainly Long Interspersed Nuclear Elements (LINEs), were activated. Full-length active LINEs and rTEs lying outside of genes were activated, suggesting that the increased transcription is not the direct result of transcription of the genes the rTEs are embedded in. We found that most activated transcription events were in proximity to KZNF binding sites, suggesting that KZNF regulation contributes to the suppression of genic and rTE transcription. Finally, we demonstrate that CCR4-NOT regulates the stability of rTE RNAs, indicating that the complex tightly controls transposon expression by repressing transcription and targeting their RNAs for decay.

## Introduction

Pervasive transcription in humans refers to the extensive activity of RNA polymerase across the genome, expressing regions not normally transcribed including intergenic regions, introns, antisense strands, and retrotransposable elements. Although a subset of these transcripts may have regulatory roles, the majority are rapidly degraded to prevent interference with gene expression, transcriptional fidelity, and genome stability. Suppressing pervasive transcription is critical for maintaining transcriptional fidelity and preventing interference with regulatory networks. Transcription from nonfunctional regions or cryptic promoters results in aberrant RNA species, disrupts chromatin organization, and interferes with core nuclear processes such as DNA replication, transcription, and RNA processing. Strong regulatory mechanisms that limit pervasive transcription and retrotransposon activity are therefore required to ensure accurate gene expression, preserve genome stability, and reduce susceptibility to transcription-associated dysfunction diseases [reviewed in (Jensen et al. 2013; Clark et al. 2011; Villa and Porrua 2023)].

LINEs (Long Interspersed Nuclear Elements) are abundant retrotransposons that influence development, genome integrity, and genome evolution(Elbarbary et al. 2016; Thompson et al. 2016; Mendez-Dorantes and Burns 2023). LINE-1 (L1) sequences comprise nearly 20% of the human and mouse genomes(Lander et al. 2001). Because L1s amplify other repeat elements, such as SINEs, L1 activity may account for as much as 50% of mammalian DNA(Graham and Boissinot 2006). Activation of LINE elements is associated with genome instability, cancer, neurological problems, and aging(Beck et al. 2011; Mendez-Dorantes and Burns 2023; Dumitrache and McKinnon 2022). LINEs and other repeat elements are silenced by Kruppel-associated box zinc-finger proteins (KZFPs), a large family of repressors(Ecco et al. 2017; Rosspopoff and Trono 2023). KZFPs suppress retrotransposable elements by recruiting the transcriptional repressor KAP1/TRIM28, which in turn recruits chromatin corepressors such as the SETDB1 histone methyltransferase, heterochromatin protein 1, and the NuRD complex(Lukic et al. 2014; Turelli et al. 2014; Fasching et al. 2015; Ecco et al. 2017; Liu et al. 2018). KZFPs also bind near promoters of genes and endogenous retrovirus LTRs to repress gene expression(Lykoskoufis et al. 2024; Ecco et al. 2017; Imbeault et al. 2017; de Tribolet-Hardy et al. 2023)Furthermore, KZFPs play an important role in suppressing pervasive transcription from rTEs and other genic regions. Given the importance of KZFPs in shaping the transcriptional landscape of genomes and the dire consequences of activating LINE elements, identifying the mechanisms of LINE silencing and the pathways that regulate KZFP activity is crucial to understanding gene expression, development, and genome integrity.

CCR4-NOT is a highly conserved complex that regulates multiple steps in gene expression in the nucleus and cytoplasm. Subunits of the Ccr4-Not complex were first identified and characterized in yeast as regulators of the TATA-binding protein and transcription initiation, but it has since been shown to regulate multiple stages of gene expression (Chalabi Hagkarim and Grand 2020; Miller and Reese 2012; Collart 2016). However, comparatively less is known about its function in metazoans, especially its nuclear functions. In the cytoplasm, CCR4-NOT functions as the major mRNA deadenylase, the rate-limiting step in mRNA decay, monitors the translatability of mRNAs, and directs the destruction of untranslatable messages and those enriched in non-optimal codons (Pavanello et al. 2023; Chen and Shyu 2011; Shirai et al. 2014; Inada and Beckmann 2024; Buschauer et al. 2020). The scaffold subunit CNOT1 is critical for bridging RNA-binding proteins and the deadenylase subunits to target mRNAs for degradation (Basquin et al. 2012; Petit et al. 2012; Zhang et al. 2022; Keskeny et al. 2019; Chen et al. 2014; Kulkarni et al. 2025). CNOT4 is a RING domain-containing E3 ligase (Collart 2016; Lau et al. 2009; Albert et al. 2002). CNOT4 is involved in the ubiquitination of the small ribosomal subunit protein eS7, which acts as a signal to pause translation during elongation as ribosomes decode non-optimal codons (Absmeier et al. 2023; Buschauer et al. 2020). Apart from its role in regulating co-translational functions, CNOT4 was recently shown to mediate the effects of codon composition on transcription in the nucleus (Garg et al. 2025).

Here, we measured ongoing transcription using pulse-labeling after rapid depletion of CNOT1 and CNOT4. Depleting either subunit caused a rapid, widespread increase in transcription across the genome, including intergenic regions. A unique set of repressed genes was identified, including the KZNF family of repressors. Consistent with the role of KZNFs in repressing retrotransposable elements, transcription of LINE and other retrotransposon elements (rTEs) increased. Thus, we have discovered a novel function for CCR4-NOT in regulating genome-wide transcription and maintaining genome integrity by suppressing pervasive transcription and transposable elements.

## Results

### Depletion of CNOT1 and CNOT4 leads to wide-spread increase in nascent transcription

Yeast Ccr4- Not regulates transcription by promoting elongation and controlling the TATA- binding protein (Reese 2013; Miller and Reese 2012; Collart 2016), but it is unclear whether human CCR4-NOT does so. We performed transient transcriptome sequencing (TT-seq) using 4-thiouracil labeling to measure ongoing transcription (Gregersen et al. 2020). DLD-1 ^TIR^ ^1^ ^+^, CNOT1 ^AID,^ and CNOT4 ^AID^ cells (Kulkarni et al. 2025) were treated with auxin for 2 or 8 h to deplete the subunits, followed by a 15-minute labeling pulse (Supplemental Fig. S1A). Immunoblotting for CNOT1 and CNOT4 confirmed rapid depletion of the proteins (Supplemental Fig. S1B). TT-seq data were normalized to a spike-in control (*S. pombe* 4-thiouracil-labeled RNA), and replicates showed very high reproducibility (Pearson’s *r* > 0.99, Supplemental Fig. S1C). Interestingly, depleting CNOT1 or CNOT4 significantly increased nascent transcription genome-wide as early as 2 h post-depletion (Fig. 1A-B). Protein depletion takes at least 1 h (Kulkarni et al. 2025), so the increase in transcription was very rapid. Thousands of differentially transcribed genes (DTGs) were identified (Supplemental Fig. S2A-D, Supplemental Table S1), of which ≥ 80% showed enhanced transcription (≥1.5 FC, p^adj^ < 0. 01 cutoff, Fig. 1C) and were predominantly protein-coding mRNAs (∼80-90% of DTGs, Fig. 1D). Furthermore, there was a significant overlap between DTGs under CNOT1 and CNOT4 depletion conditions (Supplemental Fig. S3A-B). The TT-seq data were confirmed by RT-qPCR analysis of several candidate genes, in which the majority showed increased nascent transcription when each subunit was depleted (Supplemental Fig. S4A-D). It is unlikely that altered mRNA decay is significantly affecting transcription, at least at the 2 h time point, because the response is rapid (2 h) and the median mRNA half-life in DLD-1 cells is 10.6 h (Kulkarni et al. 2025). Furthermore, our previous work showed that depleting CNOT1 (increased) and CNOT4 (decreased) had opposite effects on mRNA half-life, and yet the transcriptional response was strikingly similar (Kulkarni et al. 2025). However, the number of DTGs that met the 1.5-fold cutoff increased from 2 to 8 h of CNOT1 depletion (Fig. S2). We suspect that the increase in steady-state RNA levels of transcriptional regulators, as noted in our previous publication, accentuates the changes in transcription at later time points (Kulkarni et al. 2025) (see discussion section).

**Figure 1.**
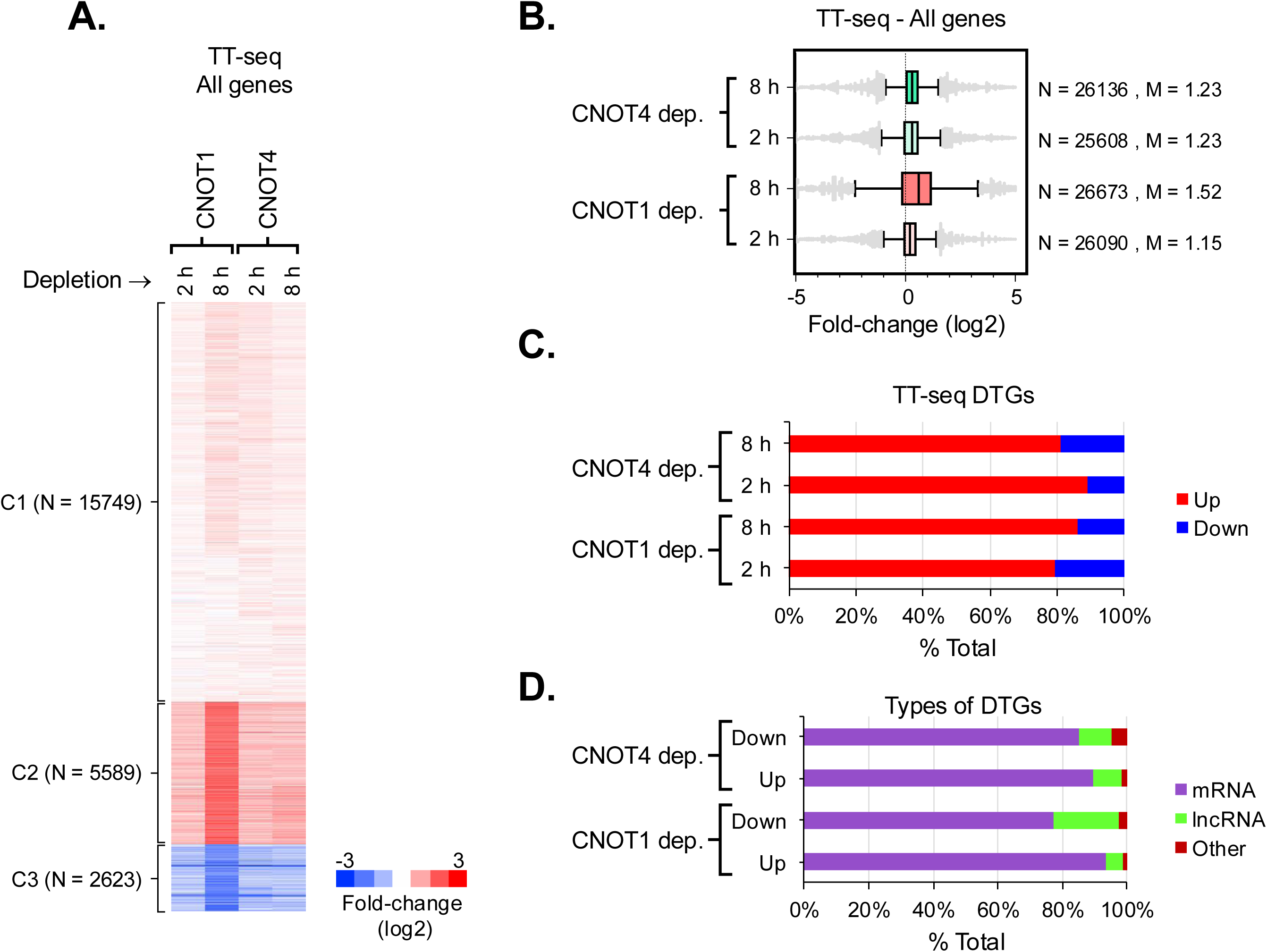
CCR4-NOT suppresses genome-wide transcription. **(A)** k-means clustering of fold-changes (FCs) in RNA transcription of genes calculated by TT-seq (N = 23961). Depleted versus undepleted [DLD-1^TIR1+^ treated with auxin] were compared. The FC in nascent transcription of genes was determined by DESeq2 analysis and normalized to a spike-in control, 4-thiouracil-labelled *S. pombe* total RNA. Heatmaps were constructed in ‘Cluster 3.0’ (K = 3 and ‘Euclidean distance’ as a similarity metric) and visualized by ‘Java TreeView’. **(B)** Boxplot analysis of the log2 FC in RNA transcription in depleted cells. The total number of datapoints (N) and median log2 fold change (M) for each group are displayed in the panel. **(C)** Frequency of up- and down regulated differentially transcribed genes (DTGs). ‘Up’ represents DTGs showing increased nascent transcription (FC ≥ 1.5, p^adj^ < 0.01) and ‘Down’ represents DTGs showing decreased nascent transcription (FC ≤ 0.66, p^adj^ < 0.01). The data are plotted as percentage of total DTGs under each depletion condition. **(D)** Distribution of the classes of DTGs. Genes were classified as mRNAs, lcRNA and all others using ShinyGO.

### CCR4-NOT regulates pervasive transcription in the intergenic regions

We next examined whether the increase in transcription is restricted to genes or also occurs in intergenic regions (IGRs). The genome was divided into 10-kb windows, and TT-seq reads from genic regions within these windows were subtracted to quantify intergenic transcription. In some cases, segments overlapping with a gene resulted in windows shorter than 10 kb. We only considered IGRs > 8 kb in our analysis. The k-means clustering plot showed a marked change in transcription within IGRs. First, under both CNOT1 and CNOT4 depletion at 2 h, increases in transcription in IGRs were more prominent than decreases (clusters C2 and C4, versus clusters C3 and C5, Fig. 2A, Supplemental Table S1). Indeed, a similar pattern was observed when we specifically analyzed differentially expressed IGRs (Fig. 2B, 2 h data points). By contrast, depletion of CNOT1 for 8 h resulted in a marked decrease in transcription across several thousand IGRs (clusters C2, C3, and C5). This pattern was underscored by the number of differentially expressed IGRs (Fig. 2B, CNOT1 - 8 h). As noted above, one possible explanation is that, by 8 h of depletion, mRNAs encoding transcriptional repressors accumulate because of the loss of deadenylase activity in CNOT1-depleted cells. This would partially restore repression of pervasive transcription (see below). Finally, we examined several intergenic loci and found evidence of pervasive transcription in IGRs in depleted cells (Fig. 2B-E). Thus, our data show that CCR4-NOT represses pervasive transcription in IGRs.

**Figure 2.**
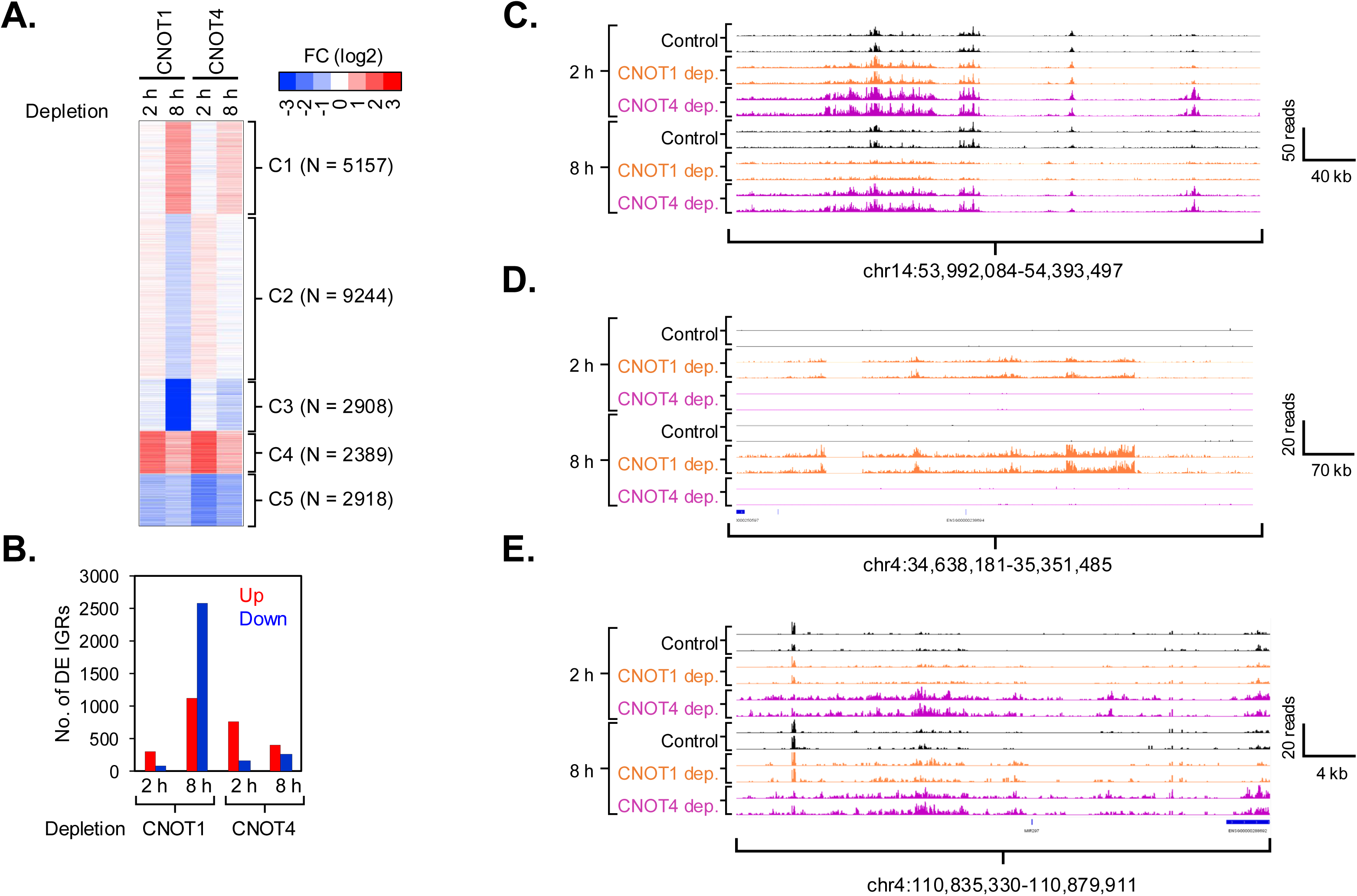
CCR4-NOT suppresses pervasive transcription in intergenic regions. **(A)** k-means clustering analysis of FC in transcription within intergenic regions (IGRs) calculated by TT-seq. FCs were calculated as described in the methods section. Each row represents an IGR window. Heatmaps were generated as described in Figure 1A. IGRs > 8kb were used in the analysis (N = 22616). **(B)** Distribution of IGRs showing significant changes in transcription under depletion conditions. ‘Up’ represents IGRs with FC ≥ 1.5, FDR < 0.05, and ‘Down’ represents IGRs with FC ≤ 0.66, FDR < 0.05. **(C-E)** IGV tracks of intergenic regions. Data from two biological replicates are displayed. Panels C, D, and E display examples where transcription was upregulated in both cells, CNOT1 only and CNOT4 only, respectively.

### CNOT1 and CNOT4 regulate transcription of the KRAB-Zinc finger repressor family genes

CNOT1- and CNOT4-depleted cells display widespread increases in genic transcription, except for genes in cluster 3, which were noticeably reduced (Fig. 1A, cluster 3). GO analysis of genes in cluster 3 revealed that ‘DNA binding transcription factor activity’ was overrepresented (Fig. 3A, 143 genes, 2.1x enrichment, 2.3E-14 FDR). Furthermore, most transcription factors in that cluster were KZNFs (Kruppel-associated box-containing zinc finger protein-coding genes), which function in gene repression (109/143, Fig. 3B). To gain further insight into this novel finding, we specifically examined ZNFs repressed under the depletion conditions and found that > 90% of the repressed ZNFs (≤ 0.67 FC, p^adj^ < 0.01) were KZNFs (Supplemental Fig. S5A). These data confirmed that although CCR4-NOT represses transcription of the majority of the genome, it positively regulates KZNF gene transcription.

**Figure 3.**
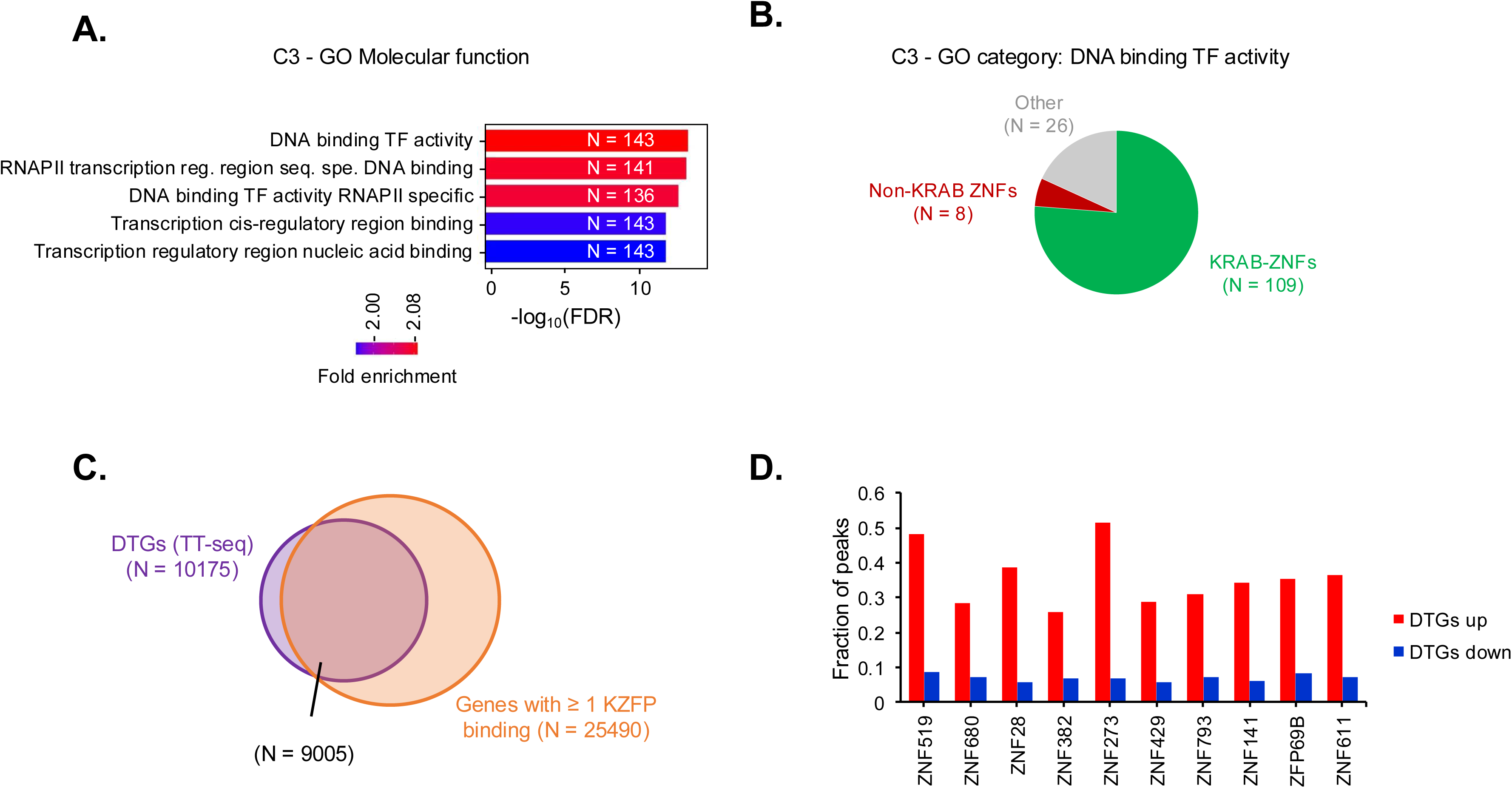
CCR4-NOT regulates KRAB Zinc Finger Protein (KZFP) expression. **(A)** Gene ontology analysis of genes in cluster 3 in Figure 1A (N = 2623). The gene set was analyzed using ShinyGO 0.80 and the top 5 GO categories based on enrichment FDR are shown. **(B)** Distribution of DNA binding transcription factor genes in the top enriched GO category in Figure 3A (N = 143). Genes encoding Zinc finger proteins (ZNFs) were grouped as with or without KRAB domain, based on(de Tribolet-Hardy et al. 2023). **(C)** Venn diagram showing overlap of DTGs and genes showing at least one KZFP binding peak. Data from KZFPs with highest number of genome-wide peaks(de Tribolet-Hardy et al. 2023) were analyzed. **(D)** Analysis of fraction of KZFP peaks in genes which are transcriptionally ‘Up’ or ‘Down’ regulated in CNOT1- and CNOT4 depleted cells (2 or 8 h depletion).

KZFPs bind throughout the genome, and we speculated that the widespread increase in nascent transcription would correlate with the presence of a KZFP binding event. We used KZFP Chip-exo data (Imbeault et al. 2017) to determine whether the KZNFs in cluster 3 (Fig. 1A) bind near genes with increased transcription. We found that ∼90% of the DTGs had at least one KZFP binding peak within or ±1KB of the gene (Fig. 3C). The binding peaks of the repressed KZFPs were more enriched at genes whose transcription increased than at those repressed by CCR4-NOT depletion (Fig. 3D). This strong association suggests that reduced KZFP expression contributes to the widespread increase in transcription in CCR4-NOT-depleted cells.

### Depletion of CNOT1 and CNOT4 increases transcription of repeat elements, particularly LINEs

KZNFs cluster on chromosome 19, having expanded and diversified in response to the rise of novel endogenous retroelements (Lupo et al. 2013; Jacobs et al. 2014; Imbeault et al. 2017; Lukic et al. 2014). Most KZNFs repressed by CCR4-NOT inactivation are on chr19, whereas KZNFs on other chromosomes typically show increased transcription, like the rest of the genome (Supplemental Fig. S5B). Examination of IGV tracks over KZNF genes on Chr19 confirms strong transcriptional repression (supplemental Fig. S5C). KZFPs bind to and repress endogenous retroviral elements and protein-coding genes by targeting promoters and ancient LTRs adjacent to genes (de Tribolet-Hardy et al. 2023; Imbeault et al. 2017; Ecco et al. 2017; Thompson et al. 2016; Chuong et al. 2017). To test whether depleting CNOT1 and CNOT4 derepresses retrotransposable elements (rTEs), the TT-seq data were remapped using the Allo tool, which efficiently allocates multi-mapped reads to genomic and offers enhanced accuracy in multimapping read assignment to repeat elements (Morrissey et al. 2024). Because this is a new tool, we compared rTE expression estimates from Allo with those from the well-established, commonly used TETranscripts program (Jin et al. 2015). Since TETranscripts does not provide quantification of individual rTE expression as Allo does, we grouped rTEs within the same rTE family to allow for comparison. The high correlation between reads for each rTE family between Allo and TETranscripts indicates that both approaches are equally effective at mapping rTEs (r = 0.98-0.99, Supplemental Fig. S6).

Differential expression analysis of rTEs mapped with the Allo tool revealed increased expression of LINE retrotransposons (LINEs) and other repeat elements, including some SINEs and LTRs of endogenous retroviruses (Fig. 4A-B, Supplemental Table S1). We then analyzed the expression of the mapped LINEs (N = 15611) and found a striking increase in LINE transcription within 2 h of depleting CNOT1 or CNOT4 (Fig. 4C, clusters 1 and 2, ∼77% of all LINEs analyzed). Furthermore, the timing of LINE activation coincided with KZNF gene repression (compare Fig. 3B to 4C). At 8 h after CNOT1 and CNOT4 depletion, global de-repression of LINE expression was still observed (Fig. 4D, clusters 2 and 4, ∼79% of all LINEs analyzed), but a subset of LINEs was re-repressed in CNOT1-depleted cells (cluster 1). The recovery of LINE repression at 8 h could be explained by increased steady-state levels of some KZNF mRNAs due to reduced mRNA decay, thereby re-establishing repression by recovering KZFP levels. When we examined the steady state levels of KZNFs in the CNOT1 depleted cells, we found an initial drop in abundance at 2hrs, but by 8 hrs the steady state levels of these messages rebounded due to enhanced stability (Supplemental Fig. S7-A-B) (Kulkarni et al. 2025). It is remarkable that compensatory effects of depletion were observed as soon as 8 hrs. This underscores the importance of examining short-term versus longer, days-long depletion of regulatory factors such as CCR4-NOT.

**Figure 4.**
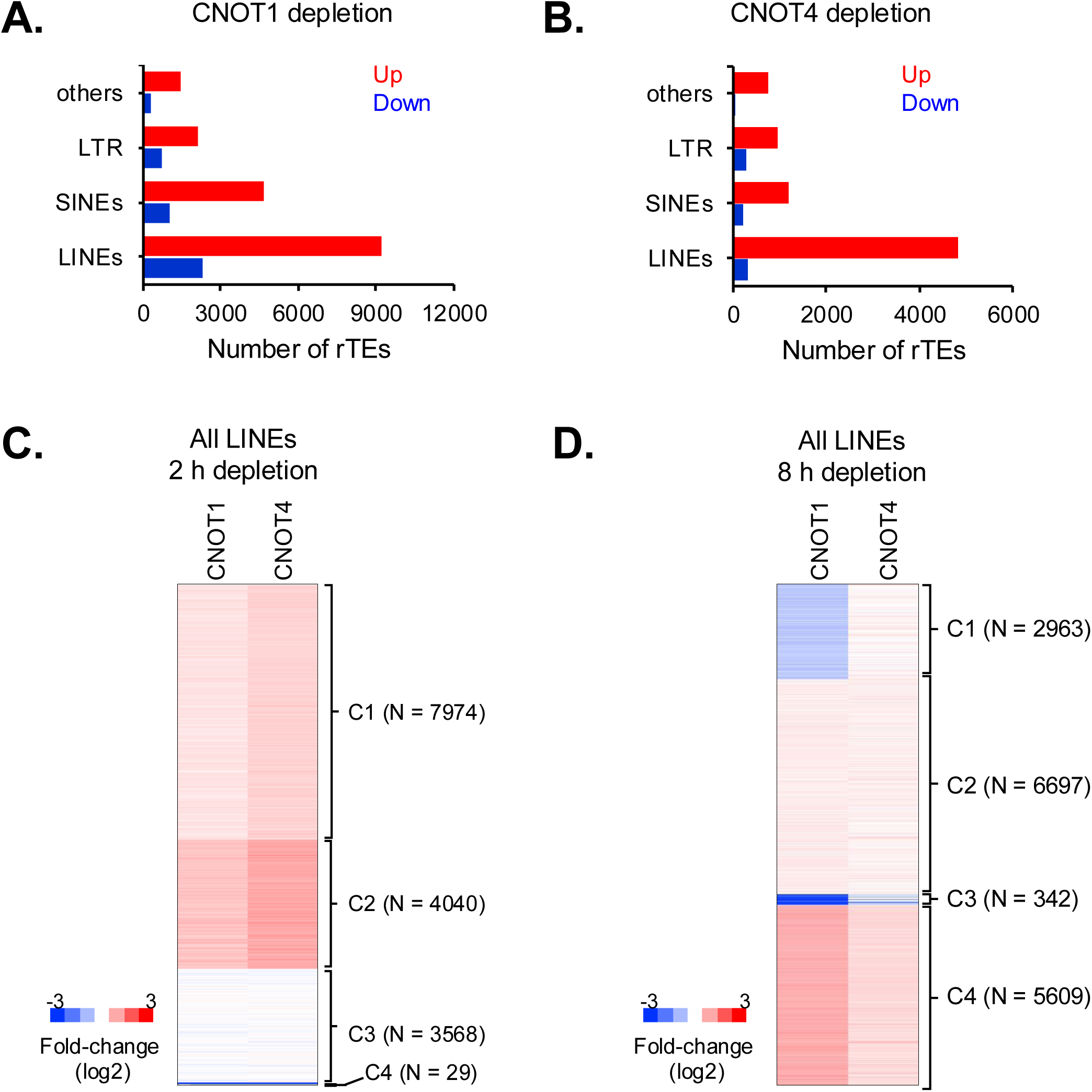
CCR4-NOT represses LINEs and other repeat transposable elements (rTEs). **(A-B)** Distribution of differentially transcribed rTEs, (FC > 1.5 and p^adj^ < 0.05) in CNOT1- **(A)** and CNOT4-depleted cells **(B)**. RNA reads were mapped using Allo, and the fold-change in transcription was determined by DESEq2. Data was normalized to the *S. pombe* spike-in control. A rTEs was scored as differentially transcribed if it met the cutoff at 2 h or 8 h depletion (cumulative). ‘Others’ include DNA transposons and other minor classes of repeat elements. **(C-D)** k-means clustering of fold-changes in transcription of LINEs after depletion of CNOT1 or CNOT4 for 2 h **(C)** or 8 h **(D)**. Heatmaps were generated as described in Figure 1A.

The majority of repeat elements, including LINEs, are located within gene bodies (≈ 90%, Supplemental Fig. S8A). To rule out the possibility that the increased reads in rTEs are mainly due to increased transcription of the gene in which they are embedded, we analyzed the transcription of rTEs located outside any annotated gene. Indeed, several hundred rTEs outside genes were also activated upon CNOT1 or CNOT4 depletion (Fig. 5A-H, Supplemental Fig. S8B-E). Functional full-length LINE-1 elements are ∼6 kb long, contain a 5′-untranslated region (5′UTR) harboring an internal promoter, two open reading frames (ORFs), and a 3′UTR (Dombroski et al. 1991). Since most LINE-1 elements are 5′ truncated and lack their promoter (Levin et al. 2025) we interrogated only those LINE-1s > 6 kb. 3% of the LINE-1s detected in our study were > 6 kb long (Fig. 6A), and several hundred of these full-length LINE-1s showed increased expression when CNOT1 or CNOT4 was depleted (Fig. 6B-C). Finally, we analyzed the putatively active LINE-1s residing in the human genome as annotated by L1Base (v2) (Penzkofer et al. 2017). Again, the majority of these active LINE-1s showed transcriptional activation under depletion conditions (Fig. 6D). Altogether, our data strongly indicate that CCR4-NOT suppresses LINE-1 expression.

**Figure 5.**
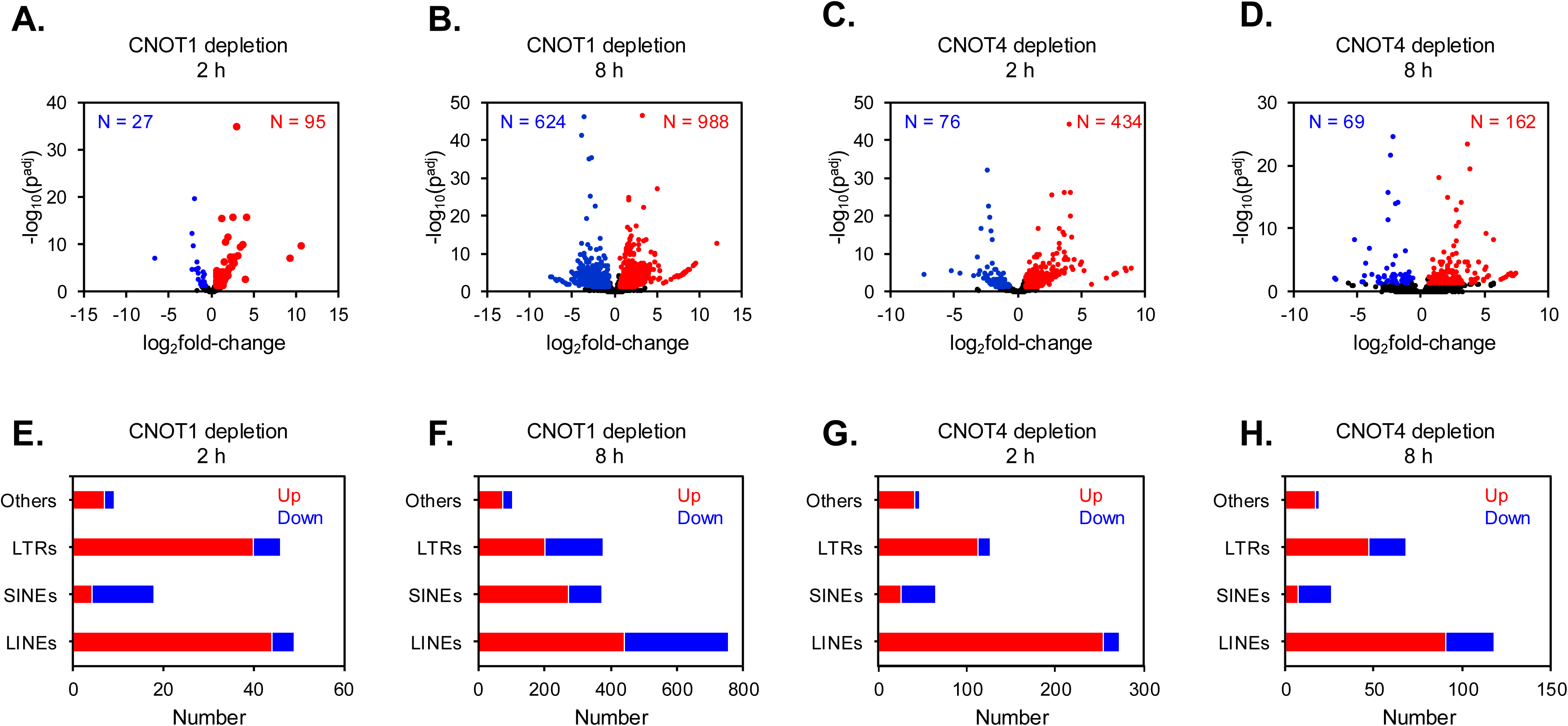
Transcription of intergenic repeat elements. **(A-D)** Volcano plots of expression of rTEs located outside of annotated genes. The number of rTEs upregulated (Up, red) and downregulated (down, blue) displaying a FC ≥ 1.5, p^adj^ < 0.05 are indicated in the panel. **(E-H)** Distribution of types of differentially transcribed rTEs from panels **A-D**.

**Figure 6.**
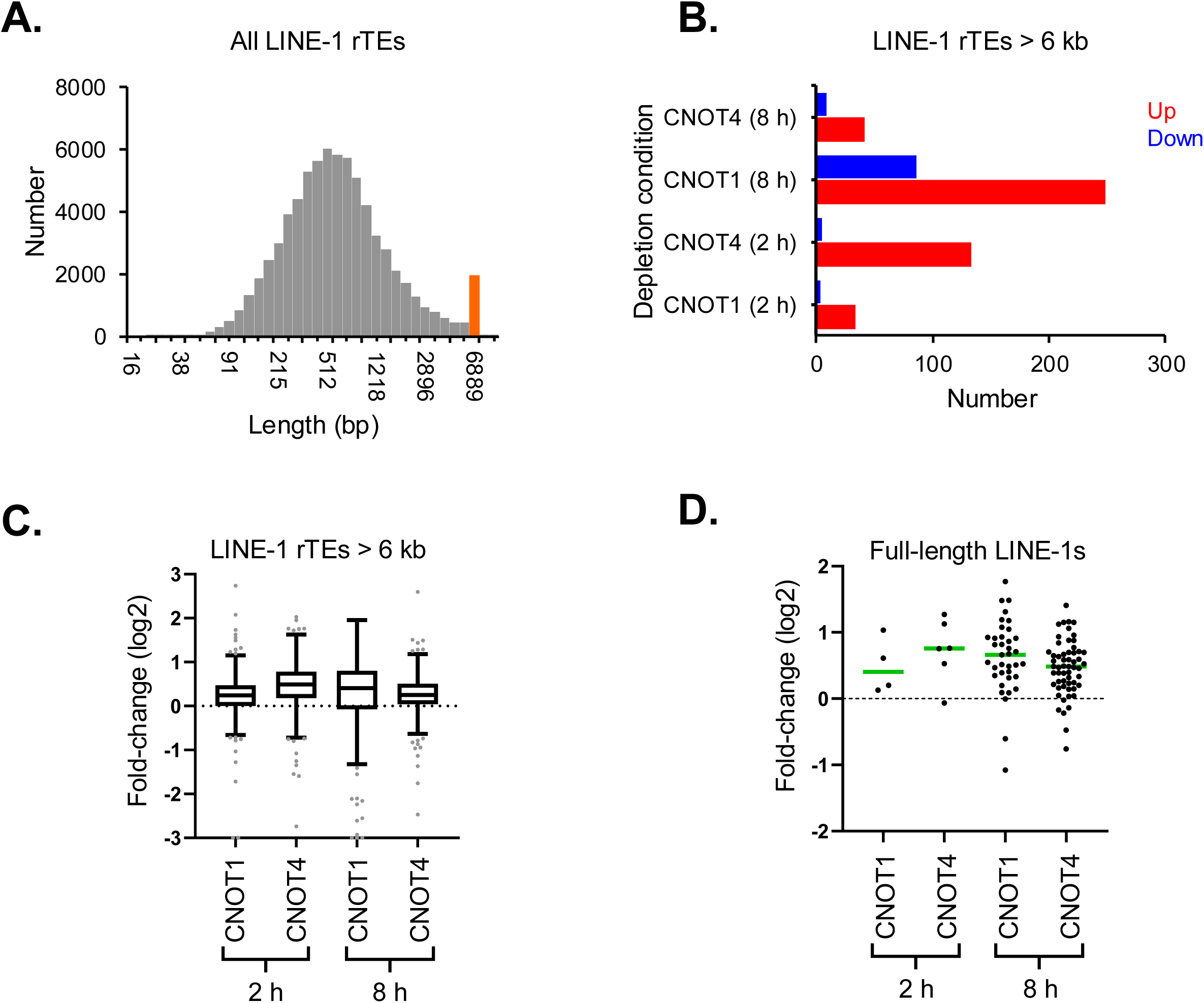
Full-length LINE-1 elements are transcriptionally activated in CNOT1- and CNOT4-depleted cells. **(A)** Length distribution of LINE-1 elements (LINE1s) mapped and detected in this study. LINE-1s > 6 kb are highlighted in orange (N = 1938). **(B)** Distribution of differentially transcribed LINE-1s > 6kb in the depletion conditions. **(C)** Boxplot analysis of fold-changes in transcription of LINE-1s > 6kb in depleted cells. **(D)** Same as in C, except full-length LINE-1s, as defined by L1Base (https://l1base.charite.de/) (Penzkofer et al. 2005) were analyzed.

LINEs influence the expression of adjacent genes by modulating chromatin, acting as enhancers, or providing alternative promoters and transcriptional start sites (TSS)(Chuong et al. 2017; Imbeault et al. 2017; Rosspopoff and Trono 2023; Thompson et al. 2016). To investigate whether changes in gene transcription are linked to LINE expression, we correlated the change in transcription of KZFP-bound LINEs with that of the adjacent gene. We focused on KZFPs that are repressed in CCR4-NOT-depleted cells and have the highest number of binding peaks (the same set of KZFPs was analyzed in Fig. 3C-D). The analysis revealed a highly significant positive correlation between changes in LINE transcription and those of their adjacent genes (Supplemental Fig. S9A-B). Similar correlations were observed with other transposable repeat elements, such as SINEs and LTRs (Spearman’s rho > 0.37, p-value < 8.42E-85, data not shown). Together, these findings suggest that CCR4-NOT-mediated gene repression is related to its functions in regulating KZFPs and LINEs.

### CCR4-NOT regulates turnover of rTE transcripts

To our knowledge, there is no comprehensive data on the half-lives of transcripts derived from rTE elements. Processed rTEs are polyadenylated, and CCR4-NOT is the major mRNA deadenylase in cells (Wiederhold and Passmore 2010). To measure the half-lives of rTE transcripts and determine whether CCR4-NOT regulates their stability, we mapped our previously published dataset of RNA expression following transcriptional inhibition with triptolide (Kulkarni et al. 2025) using Allo. We measured rTE RNA levels at different time points (0, 3, 6, and 12 h) after triptolide treatment in undepleted (DLD-1^TIR1+^) cells and in CNOT1- and CNOT4-depleted cells, and calculated their half-lives. We also separated rTEs by location (genic or intergenic) and by element type. The median half-life of rTE RNAs (8.9 h) was comparable to the median half-life of canonical RNAs (10.6 h) (Kulkarni et al. 2025). We have shown that depleting CNOT1 or CNOT4 had opposite effects on RNA half-lives. Depleting CNOT1 increased, whereas depleting CNOT4 decreased, the half-lives of RNAs. Similar to all genes, depleting CNOT1 resulted in a significant global increase in the half-lives of rTE transcripts; the median half-life increased from 8.9 h to 14.4 h (Fig. 7A). By contrast, depleting CNOT4 accelerated RNA turnover, decreasing the median half-life to 6 h. When we examined changes in rTE stability within or outside a gene, the effects of subunit depletion were observed in both groups, suggesting that CCR4-NOT regulates the turnover of rTE transcripts irrespective of their location relative to the gene body (Fig. 7B). Finally, the effects of CNOT1 and CNOT4 depletion were observed across all rTE families (Fig. 7C). Taken together, our data show that CCR4-NOT plays a role in regulating the turnover of the rTE transcriptome-wide. Thus, CCR4-NOT tightly suppresses rTEs at the transcriptional and posttranscriptional levels.

**Figure 7.**
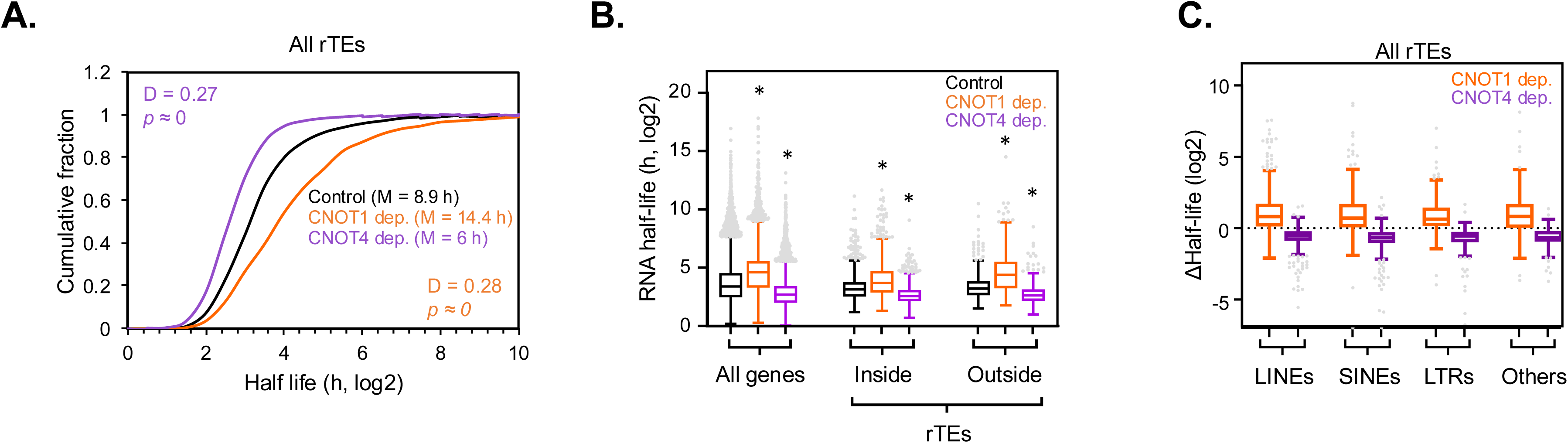
CCR4-NOT regulates turnover of rTE transcripts. **(A)** Cumulative fraction plots of half-lives of transcripts from rTEs. Half-lives were calculated by re-mapping our previously published dataset [(Kulkarni et al. 2025) and see methods] under undepleted (control, black), CNOT1- (orange) and CNOT4-depleted (purple) cells. Statistics by Two-sample Kolmogorov-Smirnov test and median half-lives in each condition (M) are shown. **(B)** Boxplots of RNA half-lives of rTE transcripts. “All genes” data include all RNAs, including mRNA and lncRNAs ([Control (N = 17869), CNOT1-depleted (N = 19216) and CNOT4-depleted (N = 20745)]. rTEs were binned based on their location with respect to annotated genes into ‘inside’ [(Control N = 934), CNOT1-depleted (N = 838) and CNOT4-depleted (N = 1431) or ‘outside’ [Control (N = 394), CNOT1-depleted (N = 316) and CNOT4-depleted (N = 565).). * *p*-value < 2.1E-43 by Mann-Whitney test. **(C)** Boxplots of changes in half-lives of rTE transcripts in depleted versus undepleted cells. Data were binned based on the rTE families and foldchanges were calculated with respect to the undepleted control.

## Discussion

CCR4-NOT has been well studied in yeast, where it was initially defined as a transcription regulator, but was later found to carry out other post-transcriptional functions [reviewed in(Chalabi Hagkarim and Grand 2020; Miller and Reese 2012; Collart 2016)]. However, much less is known about human CCR4-NOT functions in transcription. Our work fills this gap and reveals that CCR4-NOT is a global transcriptional suppressor, a function attributed, at least in part, to its regulation of KZNF expression. Using TT-seq to directly measure nascent transcription, we show that acute depletion of either CNOT1 or CNOT4 results in a rapid, genome-wide increase in transcription, affecting protein-coding genes, intergenic regions, and repeat elements. Taken together, our results suggest that, rather than acting solely as a post-transcriptional regulator, CCR4-NOT affects multiple layers of gene regulation, collectively shaping the transcriptional landscape.

Depletion of CNOT1 or CNOT4 leads to a widespread increase in nascent transcription within 2 h. This rapid response strongly argues that changes in mRNA stability cannot account for the upregulation of transcription. This conclusion is further supported by our data showing that depleting CNOT1 and CNOT4 has opposite effects on stability: CNOT1 depletion increases stability, whereas CNOT4 depletion decreases it [Fig. 7B,(Kulkarni et al. 2025)]. However, we detected additional changes in transcription after longer CNOT1 depletion (8 h), which may be augmented by changes in the steady-state levels of transcriptional regulators. For example, the transcription of a significant number of LINE elements is reduced after 8 h of CNOT1 depletion, which correlates with the recovery of KZNF steady-state mRNA levels (Supplemental Fig. 7B). Likewise, some of the other cumulative changes in genic transcription between 2 h and 8 h could result from compensatory changes in steady-state mRNA levels. On the other hand, depleting CNOT4 produces a temporally stable transcriptional response. In fact, the transcriptional state of CNOT1- and CNOT4-depleted cells is remarkably similar at 2 h (Fig. 1A, Fig. 2A, and Fig. 4C). While depleting CNOT4 accelerates mRNA turnover, the increase in transcription cancels this out, which is fully consistent with data showing that steady-state mRNA levels are relatively unchanged in CNOT4-depleted cells(Kulkarni et al. 2025). Since the steady-state levels of most transcripts did not change when CNOT4 was depleted, the enhanced mRNA turnover was compensated by increased synthesis, a process referred to as transcript buffering. Transcript buffering is a phenomenon in which a change in synthesis is balanced by an opposing compensatory change in decay to maintain steady-state RNA levels (Timmers and Tora 2018; Sun et al. 2013; Haimovich et al. 2013). Our integrated analysis of RNA synthesis, decay, and steady-state levels demonstrates that transcript buffering critically depends on CNOT1 but not CNOT4 (Supplemental Fig. S10A-B). This is explained by our previous findings that CNOT4 is dispensable for CCR4-NOT integrity and deadenylase activity. On the other hand, CNOT1 depletion disrupts CCR4-NOT and destroys deadenylase activity(Kulkarni et al. 2025; Ito et al. 2011; Gillen et al. 2021; Takahashi et al. 2020). Loss of deadenylating capacity renders CNOT1-depleted cells unable to compensate for increased transcription by ramping up decay. In contrast, CCR4-NOT and deadenylase activity are intact in CNOT4-depleted cells, enabling compensatory increases in decay that balance enhanced transcription and maintain mRNA steady-state levels.

One striking result is that, unlike the rest of the genome, transcription of KRAB zinc finger (KZNF) genes was repressed. KZFPs are the largest family of transcriptional repressors in vertebrates, and the human genome has ∼350 KZNFs; ∼200 are located within six clusters on chromosome 19 (Lukic et al. 2014). KZNFs bind ubiquitously throughout the genome, targeting promoters and binding sites within rTEs(Ecco et al. 2017; Imbeault et al. 2017; Bruno et al. 2019), and their activity would reshape the transcriptional landscape. CCR4-NOT-mediated control of KZNFs would explain the widespread increase in genic and intergenic transcription. This idea is further supported by the enrichment of KZFP binding sites near genes with increased transcription and by the strong positive correlation between transcriptional changes at KZFP-bound LINEs and adjacent genes (Fig. 3C, Supplemental Fig. 8).

KZFPs function in transcriptional silencing by repressing transposable elements (TEs), primarily retrotransposons (Ecco et al. 2017), including LINEs. LINEs are a class of non-long terminal repeat (LTR) retrotransposons with substantial potential to regulate chromatin structure and the transcription of adjacent genes (Elbarbary et al. 2016; Chuong et al. 2017; Lykoskoufis et al. 2024; Liu et al. 2018; Ecco et al. 2016). Consistent with reduced KZFP expression, we observe robust de-repression of repeat elements, most prominently LINE retrotransposons, in CCR4-NOT-depleted cells. TT-seq relies on short-read sequencing, raising the question of whether gene activation and transcription from genic promoters produce more reads over rTEs embedded in genes, or whether the loss of KZNF repression simultaneously activates both. We favor the latter. Transcription of repeat elements in intergenic regions was increased. Furthermore, transcription of full-length LINE-1 elements, including those annotated as potentially active, was upregulated. These findings place CCR4-NOT upstream of a regulatory axis linking KZFPs, rTEs, and gene expression.

The regulation of KZNFs by CCR4-NOT likely contributes to the establishment and maintenance of rTE silencing throughout cell development and growth. The rapid activation of genes and rTEs occurs within 2 h of depleting CCR4-NOT subunits, which would require the existing pool of KZNF mRNA to be reduced within this time frame. That is much faster than the median mRNA decay rate in cells (∼10 hrs). Remarkably, the steady-state level of KZNFs mRNA was significantly and strongly reduced within 2 hrs (Supplemental Fig S7). KZNF protein levels would need to drop substantially to relieve repression as well. This suggests a very elaborate and precise control over KZNF levels in cells. How precisely CCR4-NOT regulates KZNFs so rapidly remains to be determined. One possibility is that CNOT4-dependent ubiquitination targets regulatory factors that control KZFP expression, potentially via ubiquitin-dependent proteolysis or non-destructive ubiquitin signaling. This would explain why the immediate outcome (2 h). Another explanation for the rapid induction of rTEs, which would not require full depletion of KZNF protein, is the direct regulation by CCR4-NOT. CNOT3 and TRIM28 were identified as factors required for stem cell renewal and for preventing differentiation of mouse stem cells, and their promoter binding overlapped considerably (Hu et al. 2009). TRIM28/KAP1 is a corepressor utilized by KZNFs to silence rTE expression (Imbeault et al. 2017; Ecco et al. 2017). CCR4-NOT may regulate TRIM28 and repressors of rTEs.

Another significant contribution of our work is demonstrating that CCR4-NOT controls the turnover of rTE-derived transcripts. We show that depleting CNOT1 markedly stabilizes rTE RNAs across all families and genomic contexts, whereas depleting CNOT4 accelerates their decay. These opposing effects mirror those observed for canonical mRNAs (Kulkarni et al. 2025) and suggest that rTE transcripts are bona fide substrates of CCR4-NOT-mediated decay. Because rTE-derived transcripts are often polyadenylated (Borodulina et al. 2016; Perepelitsa-Belancio and Deininger 2003; Doucet et al. 2015), this 3’ end modification could render them ideal targets for CCR4-NOT-dependent RNA decay. We propose a model (Fig. 8) in which CCR4-NOT suppresses transcription of genes, intergenic regions, and repetitive elements by regulating KZNFs, while simultaneously promoting the decay of those transcripts to buffer fluctuations in transcriptional output. Such tight, redundant control mechanisms make sense given the deleterious effects of pervasive transcription and rTE activation.

**Figure 8.**
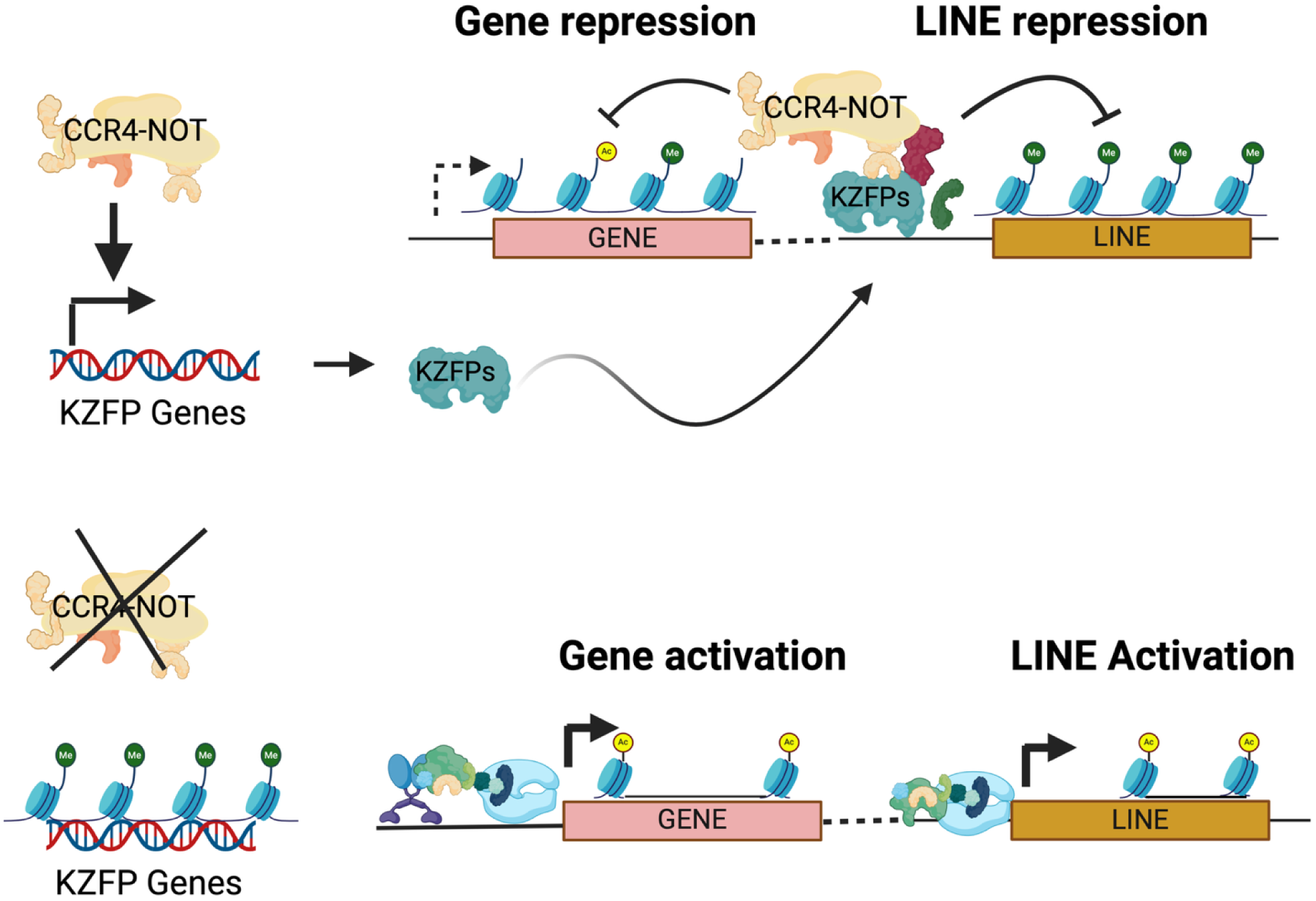
Model for the regulation of gene expression by CCR4-NOT complex. CCR4-NOT maintains normal gene expression by driving KZFP expression and directly repressing rTEs, such as LINEs. Targeting both KZNF expression and rTEs directly achieves tighter repression and provides a redundant mechanism to silence rTEs (top). Loss of CCR4-NOT reduces KZFP expression, leading to the de-repression of repeat elements and adjacent genes (bottom).

Yeast and human CCR4-NOT have been implicated in maintaining genome integrity and DNA damage responses (Jiang et al. 2019; Gaillard et al. 2009; Hagkarim et al. 2023; Chalabi Hagkarim and Grand 2020). Spurious activation of rTE, especially LINEs, causes genome instability and contributes to disease (Mendez-Dorantes and Burns 2023; Beck et al. 2011; Dumitrache and McKinnon 2022). Recently, 5-ethynyl uridine (EU) incorporation and microscopy of stained cells showed that partial CNOT1 knockdown by siRNA transfection increased bulk transcription (Hagkarim et al. 2023). This low-resolution method might have captured the increased genic and rTE transcription we demonstrated here. Interestingly, increased R-loop (RNA:DNA hybrids) formation and genome instability were also observed in the knockdown cells. R-loops are enriched in genomic regions with a higher abundance of L1 LINEs (Zeng et al. 2021). Our discovery that CCR4-NOT suppresses global transcription and rTE activation suggests a mechanism by which CCR4-NOT contributes to genome stability and integrity.

## Methods

### Cell lines

DLD-1 cells (ATCC CCL-221) were purchased from ATCC, and DLD-1^TIR1+^ cells were a kind gift from Dr. Mary Dasso at the National Institutes of Health, USA. Cells were maintained in complete media [Dulbecco’s Modified Eagle Medium (DMEM) with 10% Fetal Bovine Serum, 1X glutamax, and 1X Penicillin and Streptomycin] in a humidified incubator at 37 °C in a 5% CO2 atmosphere. Generation of AID-tagged cell lines has been described previously (Kulkarni et al. 2025).

### RNA extraction

After treatment with ±1 mM auxin for the desired time, the media were decanted, and the cells were lysed directly in 1 mL Trizol. The cells were resuspended thoroughly by pipetting up and down several times and incubated at room temperature for 5 minutes. 0.2 mL of chloroform was added per 1 mL of Trizol. The tubes were capped and shaken vigorously by hand for 15 seconds, then incubated at room temperature for 2-3 minutes. The mixture was then centrifuged at >12,000 x g for 15 min at 4 °C. The aqueous phase was collected in a separate tube, and an equal volume of 100% ethanol was added. The RNA was cleaned up using the Ambion RNA PureLink Kit according to the manufacturer’s instructions and eluted in 100 µL RNase-free water.

### Cell lysis and western blot

Cells were washed with ice-cold wash buffer (PBS supplemented with 1 mM PMSF) and harvested by scraping in wash buffer, then briefly centrifuged at high speed for 10 s. The resulting cell pellet was lysed in Laemmli buffer, boiled at 95 °C for 5–7 min, and centrifuged at high speed for 10 min. The supernatant (lysate) was collected and stored at −20 °C until analysis by Western blotting with antibodies against CNOT1 (#44613S), CNOT4 (#PA5-82214), or DDX6 (raised in-house). Proteins were detected with horseradish peroxidase–conjugated secondary antibodies and enhanced chemiluminescence (ECL), and signals were visualized with a ChemiDoc^TM^ MP imaging system.

### Transient transcriptome sequencing (TT-seq)

TT-seq libraries were generated as described previously (Gregersen et al. 2020). Briefly, cells (parental, CNOT1^AID^, and CNOT4^AID^) were grown in 150-mm dishes to reach 70% confluence at the time of auxin treatment (2 h or 8 h). Cells were then labeled by adding 4-Thiouridine (4TU, final concentration 500 µM) to the medium for 15 minutes. All work involving 4TU was performed under conditions of minimal direct light exposure. Medium was discarded, and cells were immediately lysed in Trizol. RNA extraction was carried out as described earlier. 200 µg of total RNA was mixed with 5 µg of spike-in control (total RNA from *S. pombe* labeled with 4TU, described in Supplementary Methods) in a 250-µl reaction. RNA fragmentation was carried out by adding 50 µl of NaOH (1M) to this reaction (40 minutes on ice). RNA fragmentation was stopped by adding 200 µl of Tris (1M, pH 6.8). The RNA was phenol/chloroform extracted and used for *in vitro* biotinylation using MTS-Biotin. RNA was heated at 65 °C for 10 minutes, quick-chilled on ice, and then 1X biotin buffer and MTS-biotin (Biotium) were added. Biotinylation was carried out in the dark at room temperature for 40 minutes on a rotator, followed by phenol/chloroform extraction to remove excess biotin. After heating at 65 °C and quick-chilling on ice, RNA was captured on Streptavidin beads (Dynabeads™ MyOne™ Streptavidin C1) at room temperature for 20 minutes. Beads were washed in wash buffer (100 mM Tris-HCl, pH 7.4, 1 M NaCl, 10 mM EDTA, and 0.1% Tween 20; prewarmed at 55 °C) 6 times, followed by elution by incubating in 100 mM DTT (twice, 5 minutes each). The pooled eluates were precipitated in the presence of 0.5M NaCl and 0.2 mg/ml glycogen. RNA was quantified using the Qubit RNA estimation kit.

Sequencing libraries were prepared using Illumina Stranded Total RNA Prep with Ribo-Zero Plus according to the manufacturer’s instructions. 300 ng of RNA was heated at 65 °C and quickly chilled on ice, then EPH3 buffer (from the library preparation kit) was added to adjust buffer concentrations for cDNA synthesis, followed directly by first-strand synthesis. The human (GRCh38) and *S. pombe* (ASM294v2) genomes were concatenated, and reads were mapped to the combined genome using the STAR aligner (version 2.7.11a). Raw counts of all reads mapping to the annotated region for each gene (i.e., intron + exon) were obtained using featureCounts. Gene read counts were transformed using a variance-stabilizing transformation (VST) in the DESeq2 R package. Both normalization and differential expression analysis were performed with the DESeq2 package. We used the *S. pombe* spike-in read counts for normalization. Genes showing ≥ 1.5-fold change at p-adj < 0.01 were denoted as differentially expressed.

### 4TU labelling, cDNA synthesis and RT-qPCR

4TU labeling and RNA extraction were carried out as described above. 200 µg of total RNA was used for in vitro biotinylation with MTS-Biotin. Streptavidin pull-down was performed as described above. 200 ng of purified RNA was used to synthesize cDNA with the RevertAid First Strand cDNA Synthesis Kit, according to the manufacturer’s instructions, using a random hexamer primer (prediluted 1:3). The cDNA product was used directly in PCR applications or stored at -20 °C until use. RT-qPCR was performed using PerfeCTa SYBR® Green SuperMix (Quantabio). For each condition, at least 2 experimental replicates were used. RT-qPCR was run on a CFX Opus 96 Real-Time PCR System (Biorad), and the data were analyzed using Biorad CFX Maestro 2.3 software (version 5.3.022.1030) and Microsoft Excel.

### KZFP Analysis

Raw sequencing files were downloaded from NCBI for 58 KZFPs (Supplementary Table S2) identified in cluster 3 (Fig. 1A) of our TT-seq analysis, for which binding data were previously available (Imbeault et al. 2017). Reads were trimmed with fastp v0.23.4 using default settings. Bowtie2 v2.5.1 was then used to align the single-end reads to hg38 with the “-k 25” parameter to retain multi-mapped reads. Allo v1.1.1 (Morrissey et al. 2024) was used to allocate multi-mapped reads with default settings. Peaks were called with Macs2 using default settings. Gene locations were extracted from the GENCODE v29 gtf file for hg38. A gene’s location was considered the entire gene body, including introns. The resulting bed file was intersected with the KZFP peak calls (+/- 1000 bp) using BEDTools v2.31.1.

### Analysis of transcription in intergenic regions

To assess pervasive transcription in intergenic regions, the genome was divided into non-overlapping 10 kb windows. Annotated coding regions overlapping the windows were subtracted to define intergenic regions. Reads mapping to intergenic windows were counted using featureCounts. Intergenic read counts were normalized using *S. pombe* intergenic spike-in read counts. Differential expression analysis was performed with DESeq2 to identify significantly differentially expressed intergenic windows.

### Analysis of repeat elements – TT-seq data

Reads were trimmed with fastp v0.23.4 and aligned with STAR v2.2.1 using the arguments “--outSAMmultNmax 25 --outFilterType BySJout” to retain multimapped reads. Allo v1.1.1 was used to assign multimapped reads with the arguments “--read-count --splice”. Following this, featureCounts was used to quantify transcript counts with the arguments “--countReadPairs -t exon -g gene_id -M”, which retains multimapped reads. The annotation file used in featureCounts was constructed from the GENCODE v29 gtf file and the annotated repeats from RepeatMasker v4.1.3 for hg38. After read summarization, DEseq2 was used to analyze differentially expressed transposable elements. In this analysis, the scaling factors calculated from the spike-in described above were used for normalization. Additionally, canonical genes were removed from the table before DE analysis. We retained them during the featureCounts step to avoid transposable elements that overlapped gene exons.

### Calculation of half-lives of rTEs

To calculate RNA half-lives of rTEs, previously published sequencing data from our lab were interrogated (Kulkarni et al. 2025). Briefly, DLD-1^TIR1+^, CNOT1^AID^, and CNOT4^AID^ cells were treated with auxin (1 mM), followed by triptolide (1 μM) for 0, 3, 6, and 12 h (in the presence of 1 mM auxin). Reads from the RNA-seq data were mapped using the Allo tool (Morrissey et al. 2024) and fold changes in expression relative to the 0 h treatment were calculated using DESeq2 analysis (with ERCC spike-in normalization). The decay rate constant (k_decay_) was then calculated, followed by half-life determination [t_1/2_ = ln(2)/k_decay_].

### Reagents

All the relevant reagents used in this study are listed in Supplementary Table S3.

### Webtools and softwares

All the webtools and softwares used in this study are listed in Supplementary Table S4.

## Data availability

All primary data obtained from TT-seq have been deposited at the Gene Expression Omnibus (Accession ID: GSE308749).

## Author Contributions

S.K. performed the molecular and cell biology experiments. S.K., A.M., A.S., O.T.A., I.A. and S.M. performed gene mapping and bioinformatics analysis. C.A.K. performed the sequencing. S.K. and J.C.R. conceived the experiments and wrote the paper.

## Acknowledgements

We thank members of Reese lab and that of the Penn State Center for Eukaryotic Gene Regulation for their helpful comments and suggestions. We would also like to thank core facilities at the Penn State University including Genomics (RRID:SCR_023645) and Genomics Research Incubator (RRID:SCR_024530). Mary Dasso (NIH) is recognized for providing DLD-1 Tir1 expressing cells and plasmids. We also acknowledge Siqing Wang (CHOP Research Institute) for their advice on TT-seq library preparation. This research was supported by funds from National Institutes of Health R35 GM136353 (J.C.R) and R35 GM144135 (S.M.).

