## supplemental information for "Human CCR4-NOT suppresses pervasive transcription and retrotransposable elements"

### Supplemental Fig. S1

A.

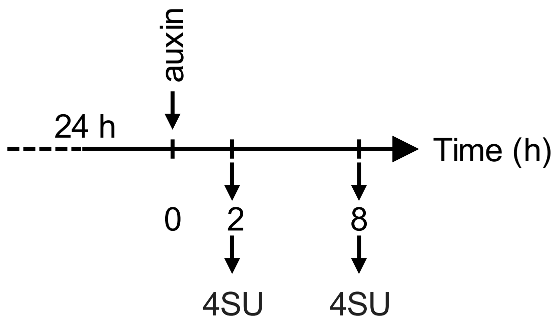


B.

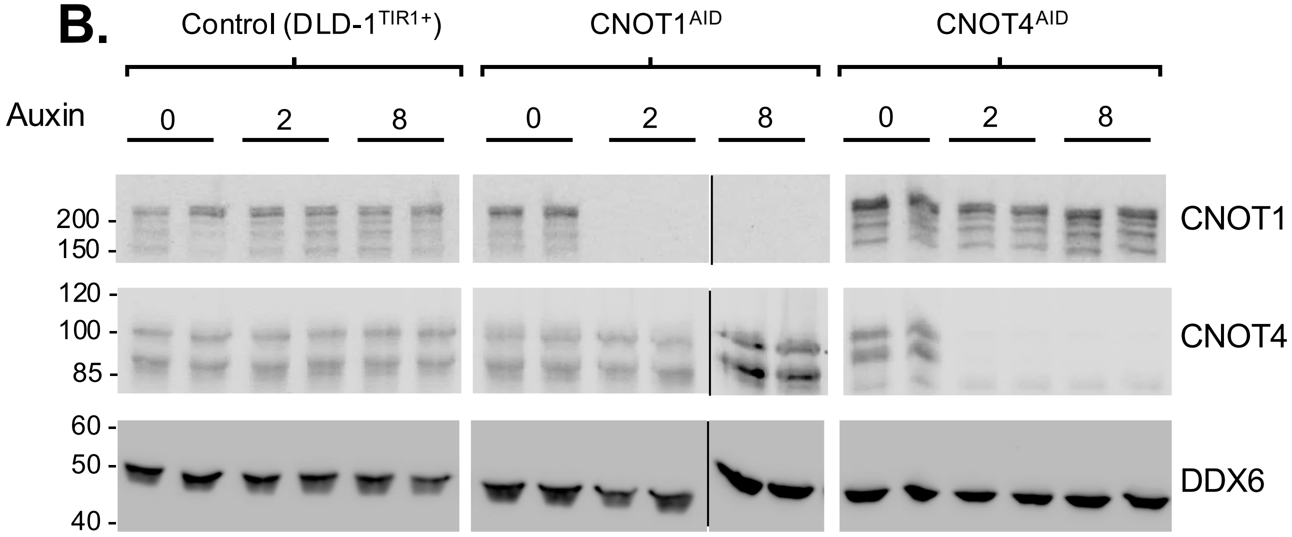


C.

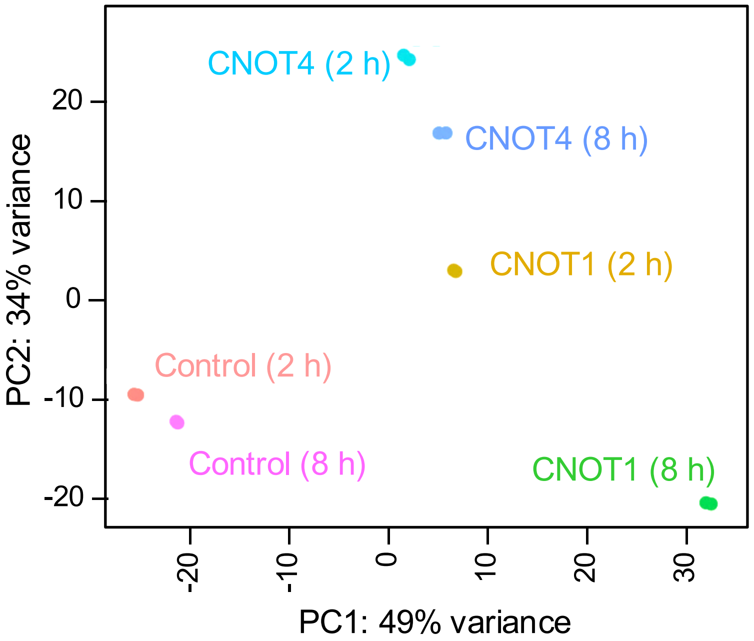

### Supplemental Fig. S2

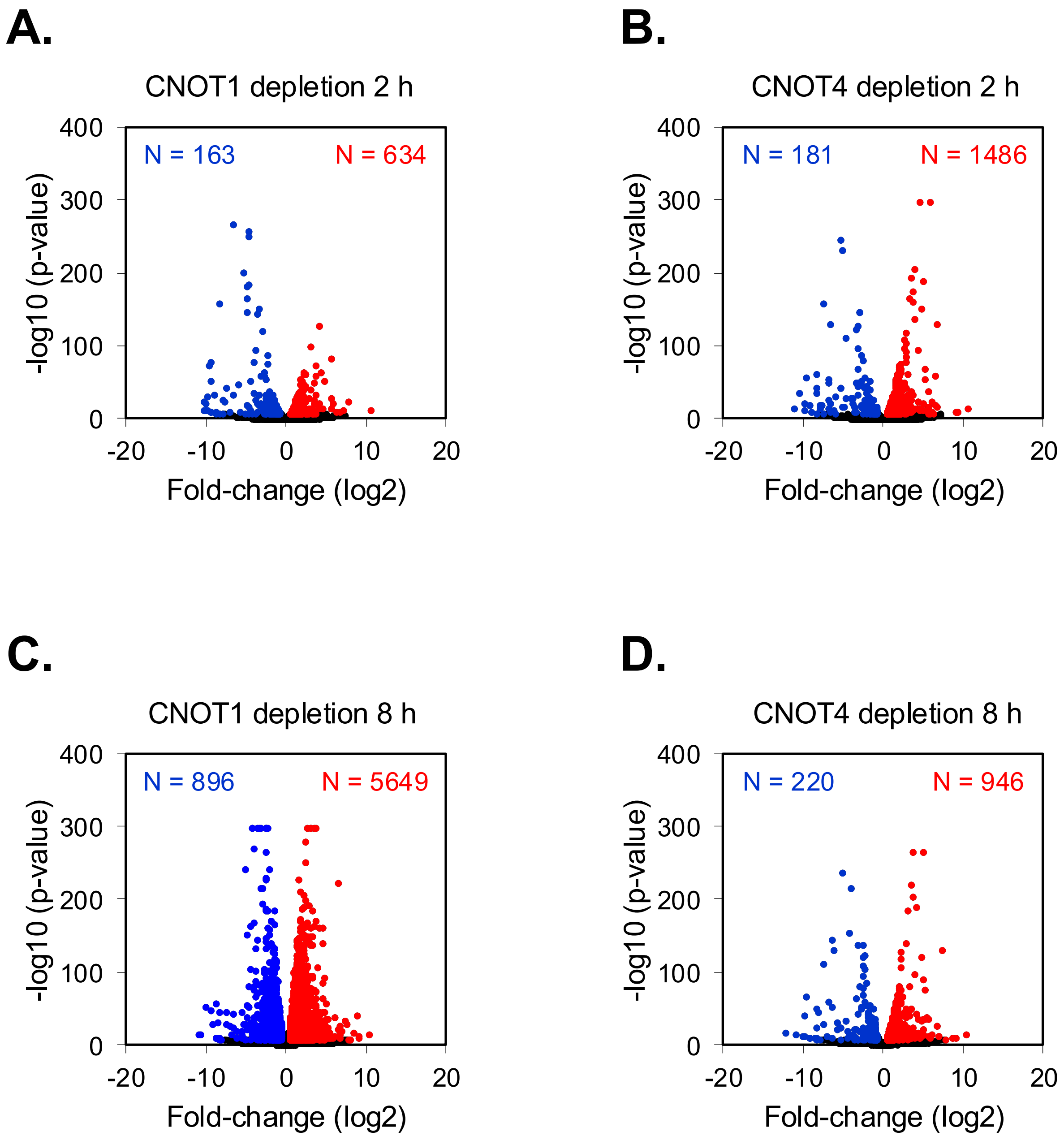

### Supplemental Fig. S3

A.

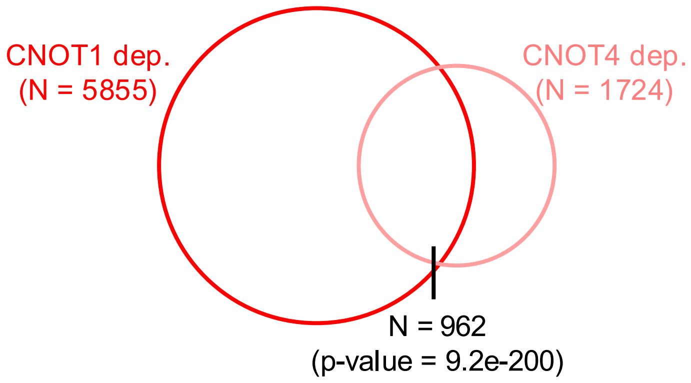


B.

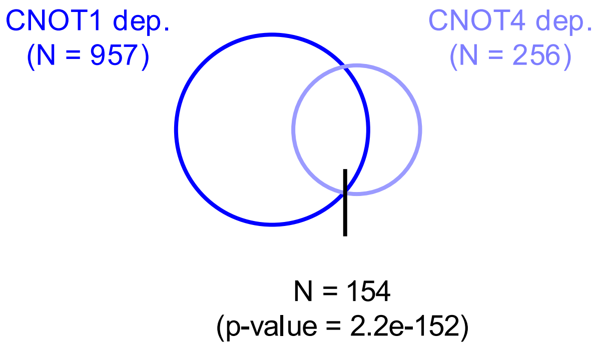

### Supplemental Fig. S4

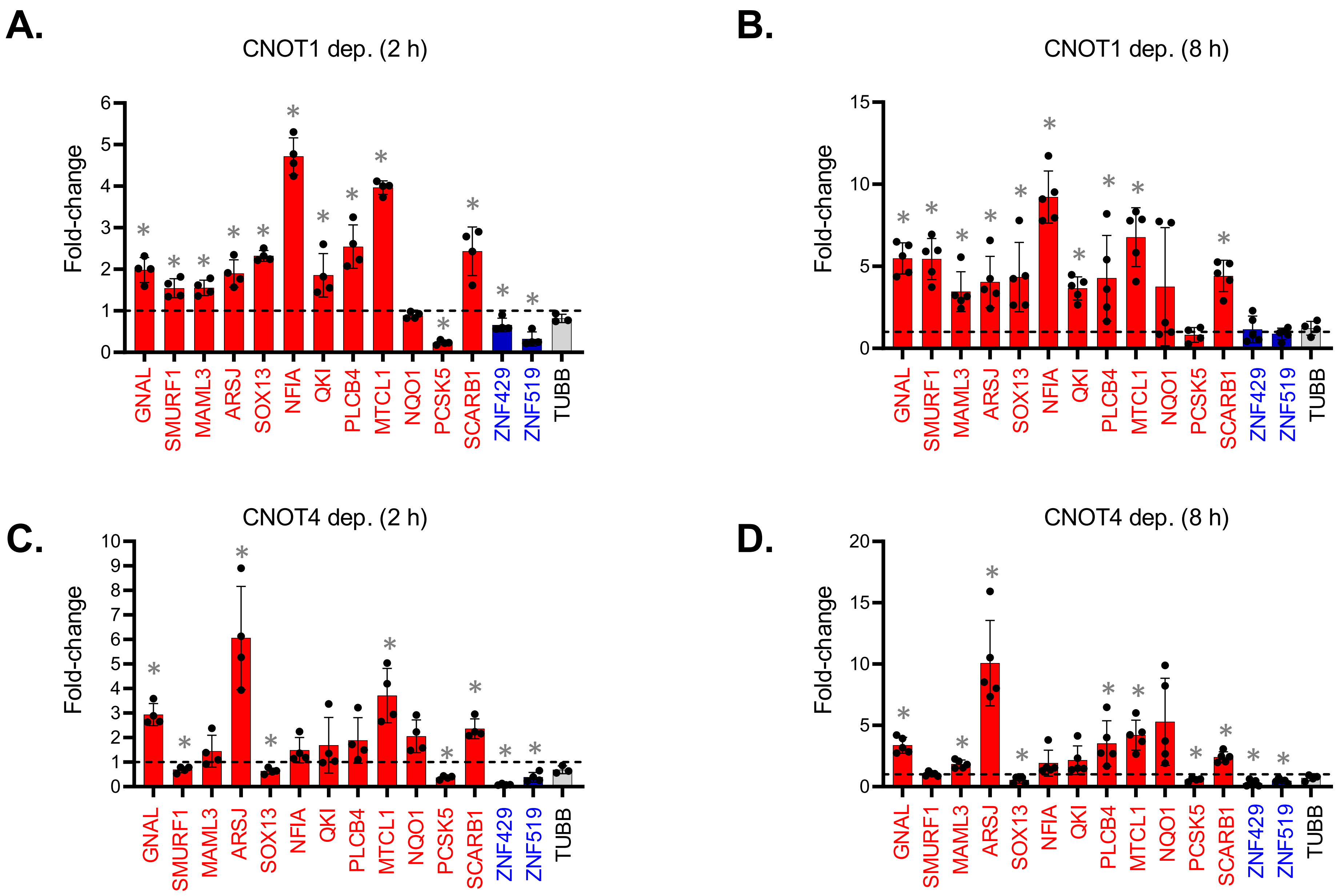

### Supplemental Fig. S5

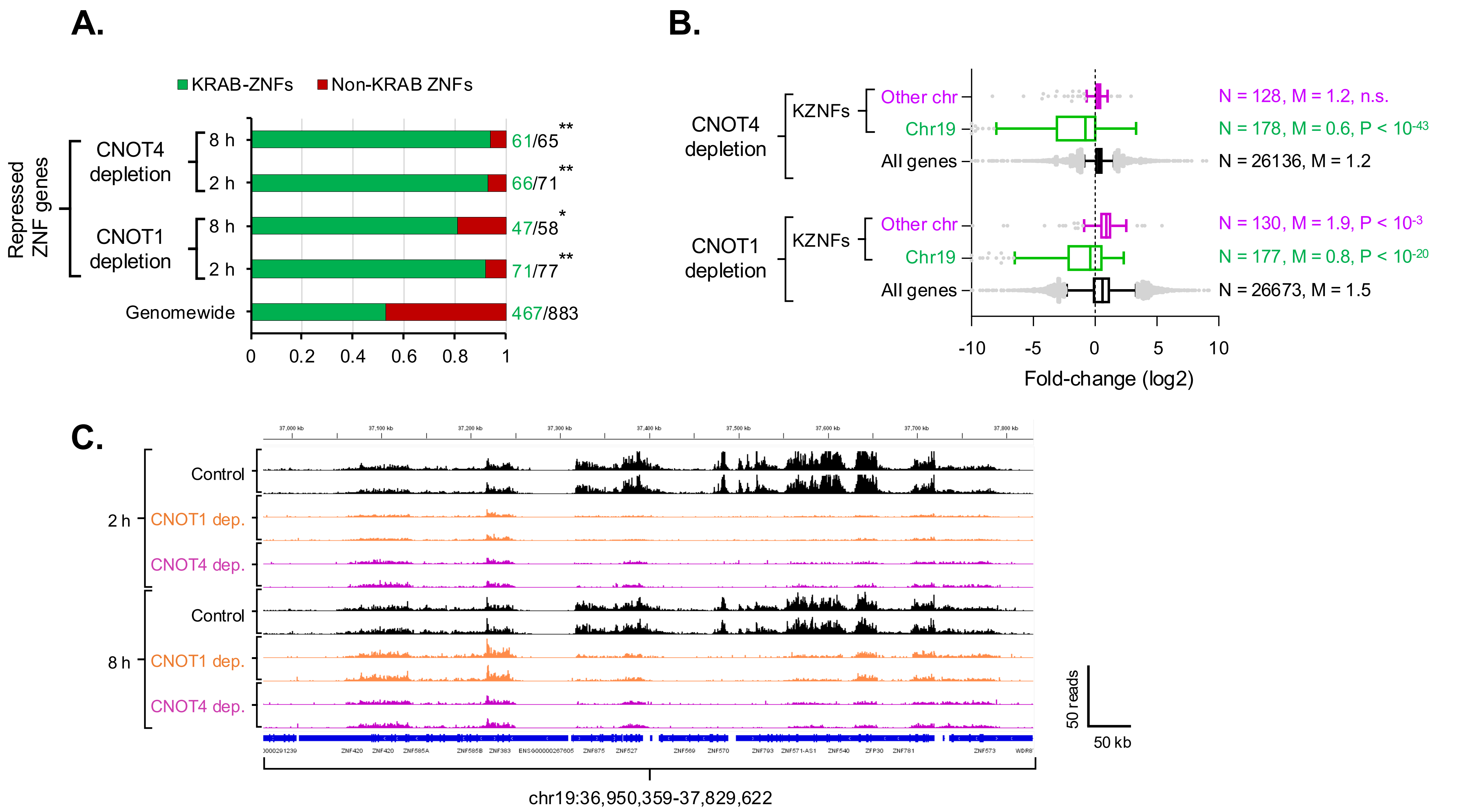

### Supplemental Fig. S6

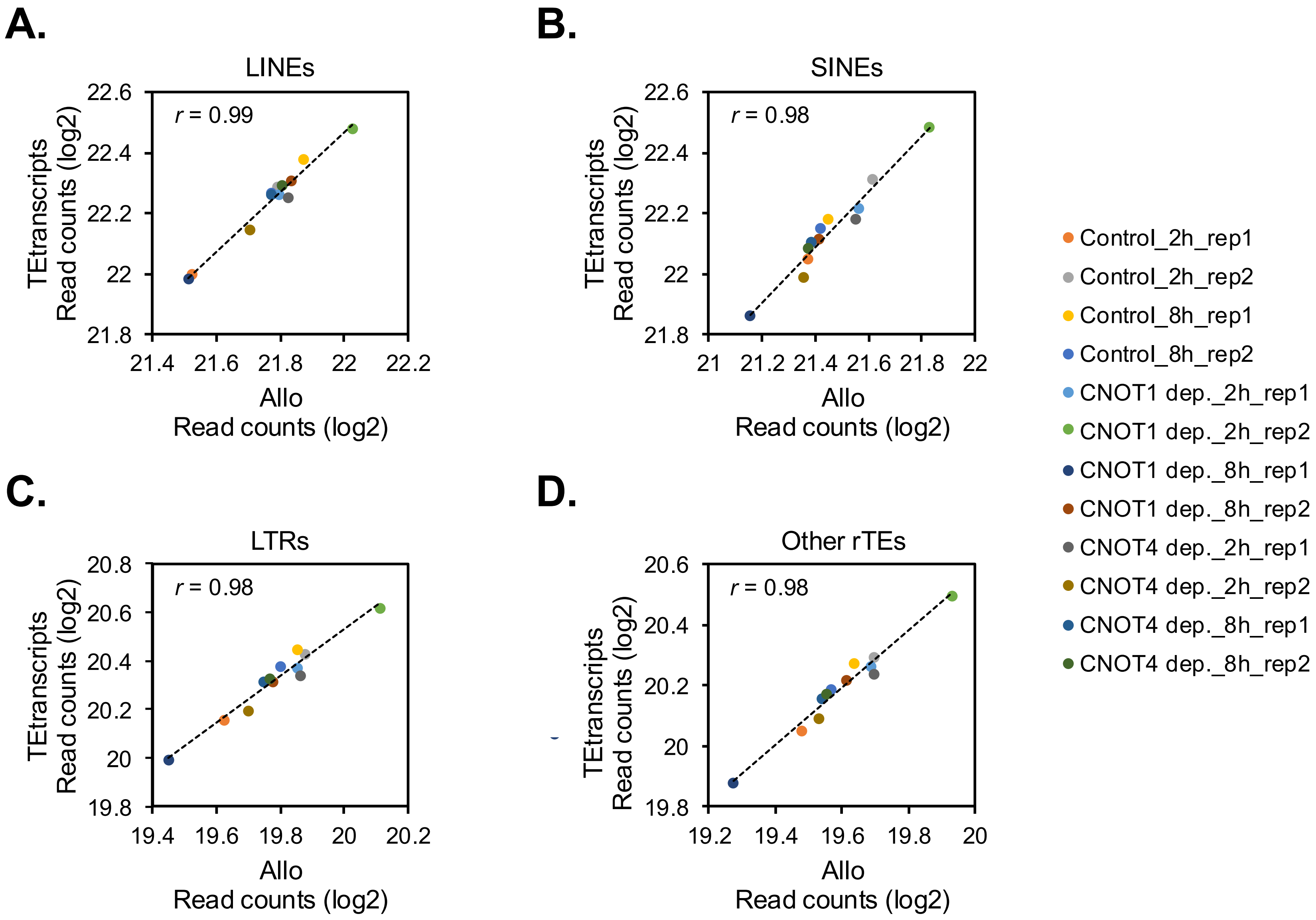

### Supplemental Fig. S7

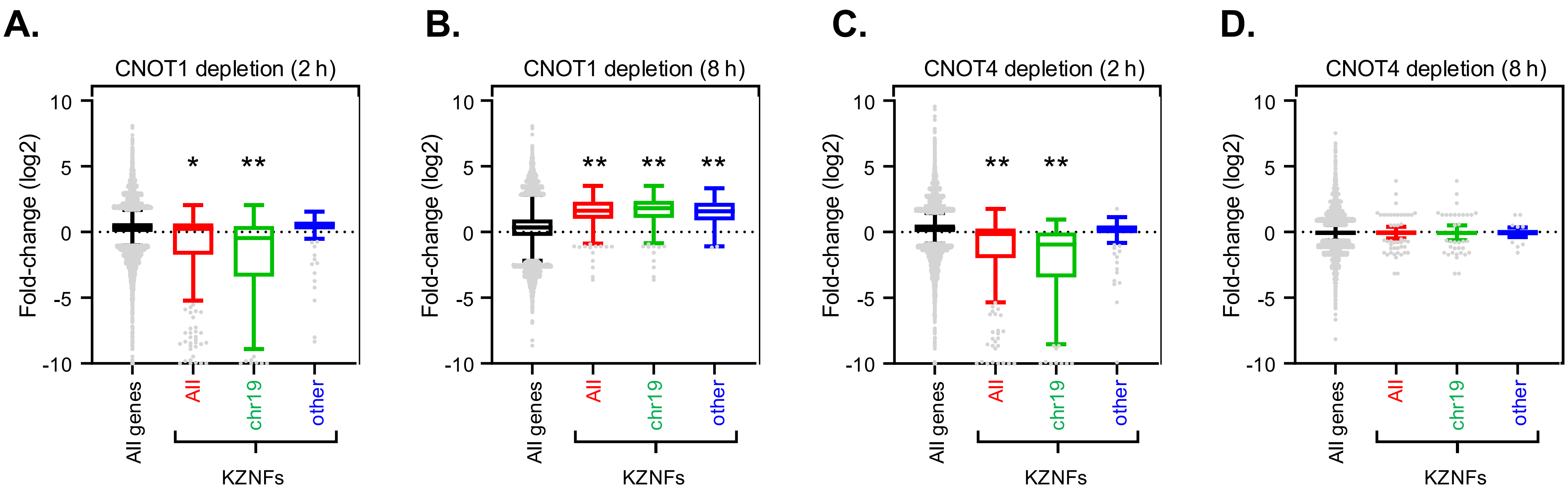

### Supplemental Fig. S8

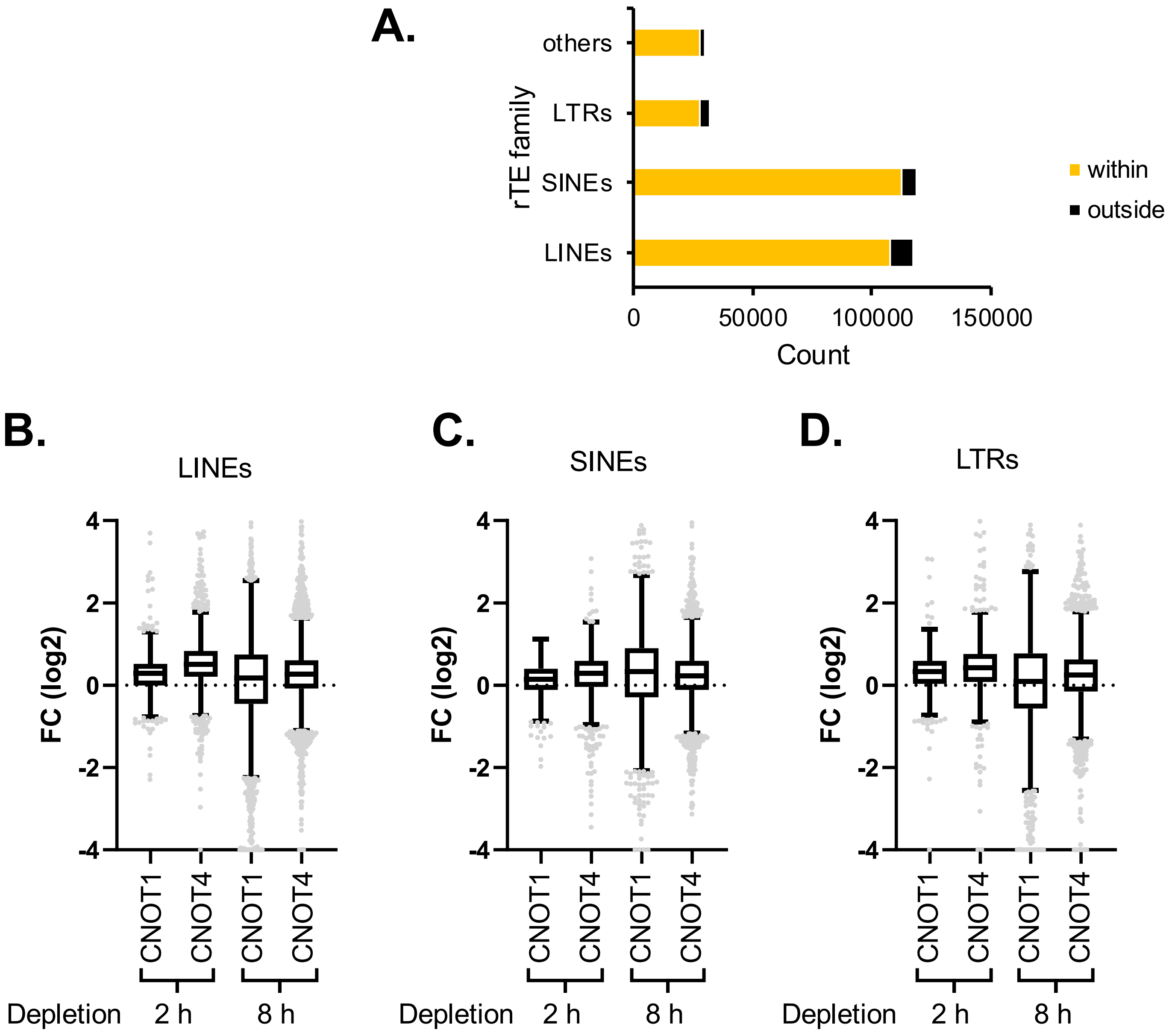

### Supplemental Fig. S9

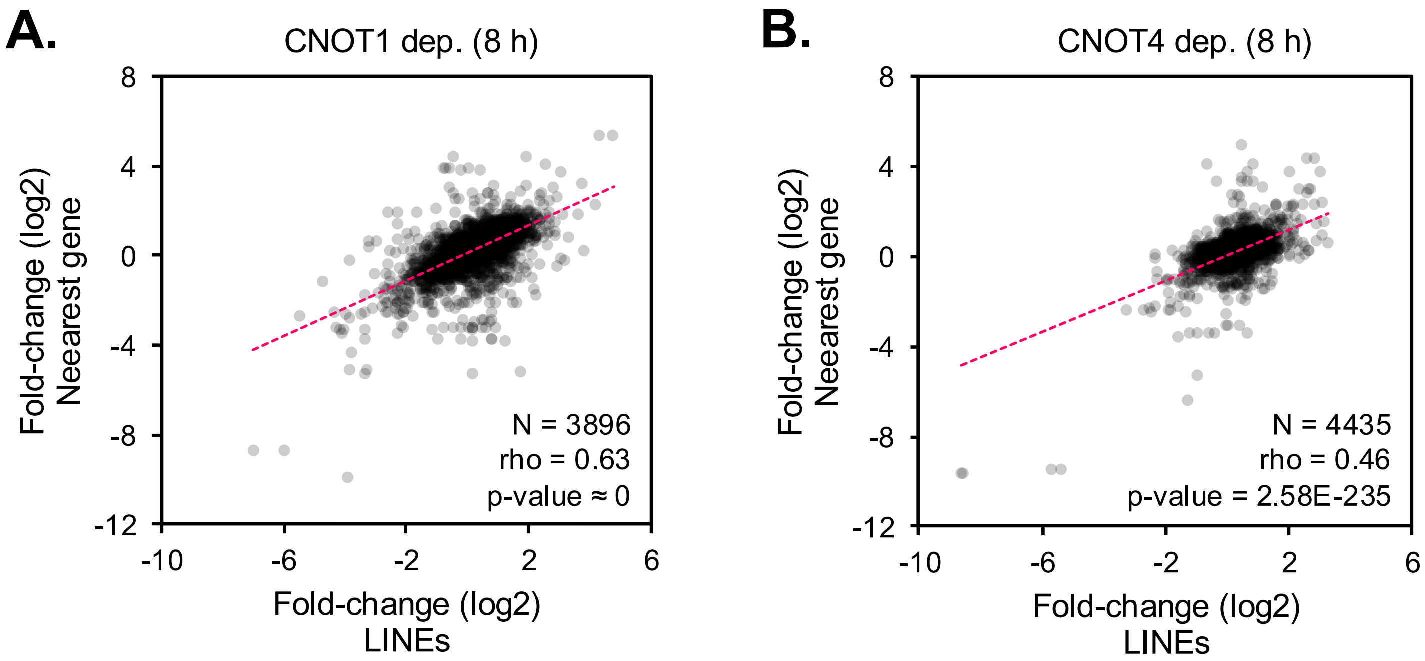

### Supplemental Fig. S10

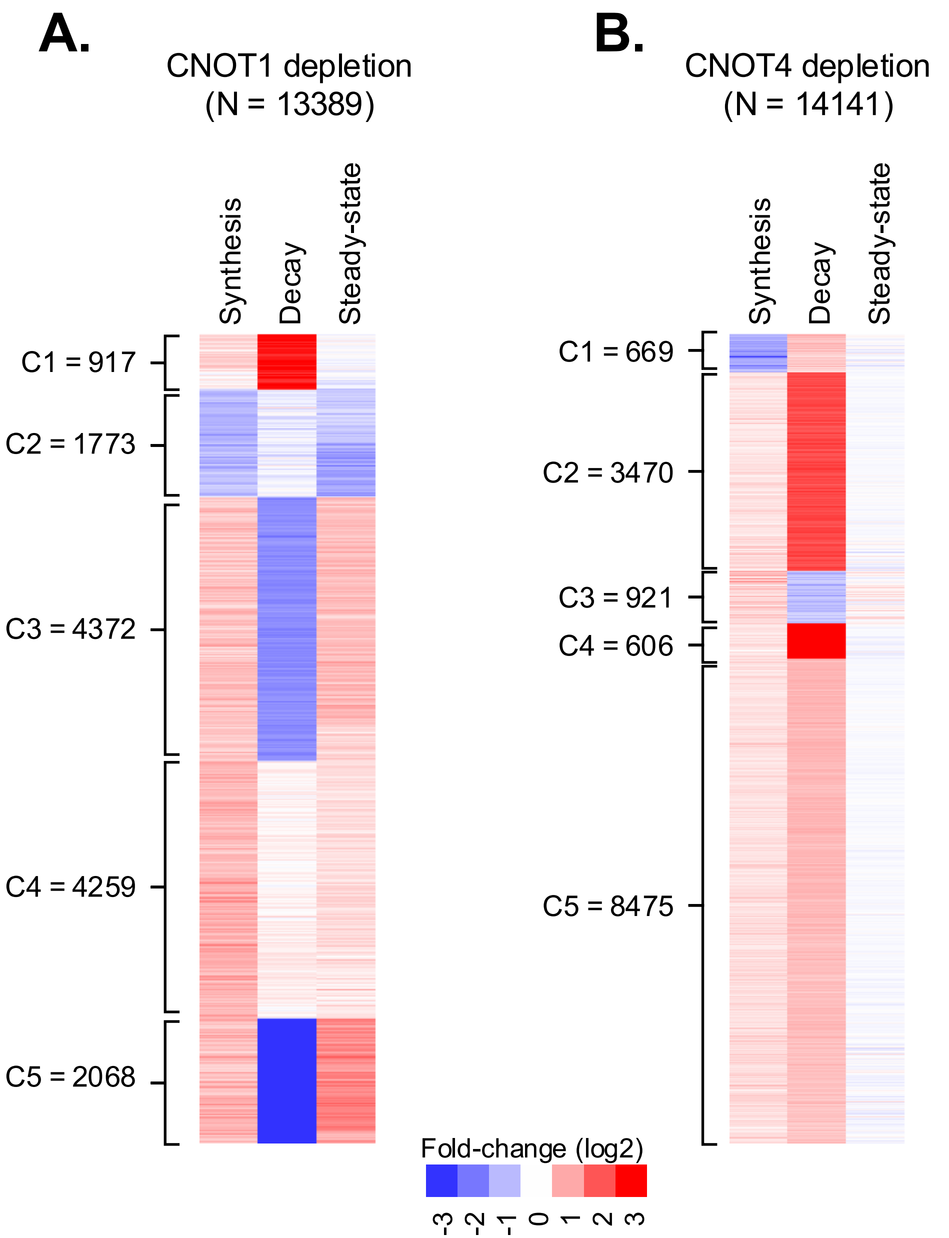

**Supplementary figures legends**

**Supplemental Fig. S1. Supporting data for TT-seq experiment.** (A) Schematic depicting the experimental setup to study nascent transcription by transient transcriptome sequencing (TT-seq). (B) Immunoblot analysis for CNOT1 and CNOT1 after 2 and 8 h auxin treatment. Lysates from control (DLD-$1^{TIR1+}$), $CNOT1^{AID,}$ and $CNOT4^{AID}$ cells treated with 1 mM auxin for 0, 2, and 8 h were immunoblotted for CNOT1, CNOT4, and DDX6 (loading control). Lysates from two biological replicates of each timepoint were analyzed. The vertical line indicates the splicing of two blots that were run in parallel. Molecular weight markers in kDa are shown on the left. The increase in CNOT4 protein levels in CNOT1-depleted cells (8 h) results from increased CNOT4 mRNA levels, as described in our published work (Kulkarni et al. 2025). (C) Principal component analysis of TT-seq data. The two biological replicates for each condition were plotted.

**Supplemental Fig. S2. Depletion of CNOT1 and CNOT4 results in genome-wide transcriptional activation.** (A-D). Volcano-plots of $\log_2$FoldChange (FC) in nascent transcription versus negative $\log_{10}$ of p-values based on DESeq2 analysis. The FCs (depleted versus undepleted) were determined by DESeq2 and normalized to a spike-in (4-thiouracil labelled *S. pombe* total RNA). The number of differentially transcribed genes (DTGs) with an FC ≥ 1.5 and $p^{adj} < 0.01$ is indicated in each panel.

**Supplemental Fig. S3. Analysis of DTGs identified using TT-seq.** (A) Venn diagram showing significant overlap between transcriptionally up-regulated genes (2 or 8 h of depletion). P-value of overlap was calculated by R-studio. (B) Same as in A, except transcriptionally down-regulated genes were analyzed.

**Supplemental Fig. S4. Validation of TT-seq results by candidate gene analysis by RT-qPCR.** (A-D). RNA isolated from 4-thiouracil labeled cells was biotinylated, captured on streptavidin beads, eluted and subjected to RT-qPCR. Expression was normalized to 18S rRNA expression. Fold-changes with respect to the undepleted control set to 1.0 (DLD-$1^{TIR1+}$ treated with auxin) are plotted. Data represent average ± S.D. from ≥ 3 experimental replicates, ∗ indicates $p < 0.05$ determined by unpaired two-tailed Student's t test. Genes highlighted in red and blue indicate transcriptionally activated and repressed genes, respectively, as defined by TT-seq.

**Supplemental Fig. S5. CCR4-NOT regulates KZNF expression.** (A) Analysis of genes coding for zinc finger proteins whose transcription is significantly repressed in depletion conditions. Data shows the fraction of these genes with or without KRAB domain according to(de Tribolet-Hardy et al. 2023). P-values of the over-representation compared to the genome-wide fraction of ZNF versus KZNFs were calculated by Fisher's exact test (* two-tailed p-value < 0.05, ** two-tailed p-value < 0.005). (B) KZNFs on chromosome 19 are repressed in depleted cells. Boxplot analysis of changes in transcription of KRAB-ZNFs (KZNFs) genes after 8 h depletion of CNOT1 or CNOT4. KZNFs were binned based on their location on chromosome 19 or other chromosomes (de Tribolet-Hardy et al. 2023). Total number of datapoints (N), median fold-change (M) and P-values by Mann-Whitney Test (compared to all) are indicated in the panel. (C) IGV tracks of KZNF genes on chromosome 19. Data from TT-seq experiment were visualized using Integrative Genomics Viewer (version 2.14.1). The track heights were adjusted according to *S. pombe* reads, which were used as a spike-in control in the TT-seq experiment. Data from 2 biological replicates were visualized.

**Supplemental Fig. S6. Comparison of the mapping by Allo and TEtranscripts** (A-D) Comparison of the numbers of TT-seq read counts of rTEs mapped using Allo tool (Morrissey et al. 2024) versus TEtranscript(Jin et al. 2015). The data were binned into LINEs (A), SINEs (B), LTRs (C), and other minor repeat classes (D). Pearson's correlation coefficients (*r*) are shown in each panel.

**Supplemental Fig. S7. Analysis of steady state mRNA levels under depletion of CNOT1 and CNOT4.** (A-B) Our previously published datasets (Kulkarni et al. 2025) of steady-state mRNA were reanalyzed to quantify fold changes in KZNFs compared to all genes in CNOT1- and CNOT4-depleted cells. 'All genes' represent all genes whose changes in steady-state RNA levels can be calculated (N ≥ 22489). KZNFs were binned based on their location on chromosome 19 or other chromosomes (de Tribolet-Hardy et al. 2023). 'All KZNFs' (N ≥ 306); 'chr19-KZNFs' (N ≥ 173); 'KZNFs-other' (N ≥ 133). P-values by Mann-Whitney Test compared to 'All genes' (* $p < 1.0E-09$, ** $p < 1.0E-24$).

**Supplemental Fig. S8. rTEs located outside annotated gene sequences are transcriptionally activated under depletion conditions.** (A). Distribution of rTEs based on their location with respect to an annotated gene. rTEs were separated into those within or outside the gene. (B-E) Boxplot analysis of changes in transcription of rTEs located outside of an annotated gene. Data from TT-seq were plotted for LINEs (B), SINEs (C), LTRs (D) and other minor classes of rTEs (E).

**Supplemental Fig. S9: Transcription of LINEs correlates with adjacent gene expression.** (A-B). Correlation plots between fold-change in transcription (TT-seq) of LINE elements versus that of its nearest gene after 8 h of depletion of CNOT1 (A) or CNOT4 (B). The top ten KZFPs with the highest number of genome-wide peaks determined by ChIP-exo (Imbeault et al. 2017) and whose transcription was repressed upon CNOT1 or CNOT4 depletion were analyzed (as in Figure 7A-B). LINEs associated with these KZFPs were mapped, and the nearest gene was located for each LINE, followed by correlation analysis. Number of datapoints (N), Spearman correlation (rho) and 2-sided p-value are shown on each plot.

**Supplemental Fig. S10. Comparison of the change in synthesis and decay versus steady state RNA in depleted cells.** (A-B) k-means clustering of fold-changes in RNA synthesis, RNA decay and steady-state RNA for all genes. Foldchanges in RNA synthesis upon 8 h of depletion of CNOT1 (A) or CNOT4 (B) were calculated using TT-seq data (8 h depletion, this study), while changes in RNA decay and steady-state RNA levels (8 h depletion) under depletion conditions were calculated from our previously published dataset(Kulkarni et al. 2025). The rows represent individual RNAs. FC data were used to make heatmaps in 'Cluster 3.0' (K = 5 and 'Euclidean distance' as a similarity metric) and were visualized by 'Java TreeView'.
